# Distinct expression of select and transcriptome-wide isolated 3’UTRs suggests critical roles in development and transition states

**DOI:** 10.1101/2020.12.11.420125

**Authors:** Shaoyi Ji, Ze Yang, Leonardi Gozali, Thomas Kenney, Arif Kocabas, Carolyn Jinsook Park, Mary Hynes

**Author notes:** Corresponding author Mary Hynes 327 Campus Drive, Dept of Biology, Stanford University. Co-first authors (Shaoyi Ji, Ze Yang). **Author Contributions:** Shaoyi Ji and Ze Yang contributed equally to the manuscript; Dr. Ji initiated and carried out all bioinformatics analyses. Dr. Yang initiated all work on neural and hair cell progenitors and coordinated and oversaw the work of research assistants (Leonardi Gozali and Thomas Kenney) who carried out many of the ISH studies. Dr. Kocabas initiated work on the CYTOO chips overseeing research assistants Saranya Santoosh Kumar and Carolyn Park. **Competing Interest Statement:** No competing interests.

## Abstract

Mature mRNA molecules are typically considered to be comprised of a 5’UTR, a 3’UTR and a coding region (CDS), all attached until degradation. Unexpectedly, however, there have been multiple recent reports of widespread differential expression of mRNA 3’UTRs and their cognate coding regions, resulting in the expression of isolated 3’UTRs (i3’UTRs); these i3’UTRs can be highly expressed, often in reciprocal patterns to their cognate CDS. Similar to the role of other lncRNAs, isolated 3’UTRs are likely to play an important role in gene regulation but little is known about the contexts in which they are deployed. To begin to parse the functions of i3’UTRs, here we carry out *in vitro, in vivo* and *in silico* analyses of differential 3’UTR/CDS mRNA ratio usage across tissues, development and cell state changes both for a select list of developmentally important genes as well as through unbiased transcriptome-wide analyses. Across two developmental paradigms we find a distinct switch from high i3’UTR expression of stem cell related genes in proliferating cells compared to newly differentiated cells. Our unbiased transcriptome analysis across multiple gene sets shows that regardless of tissue, genes with high 3’UTR to CDS ratios belong predominantly to gene ontology categories related to cell-type specific functions while in contrast, the gene ontology categories of genes with low 3’UTR to CDS ratios are similar and relate to common cellular functions. In addition to these specific findings our data provide critical information from which detailed hypotheses for individual i3’UTRs can be tested-with a common theme that i3’UTRs appear poised to regulate cell-specific gene expression and state.

**Significance Statement:** The widespread existence and expression of mRNA 3’ untranslated sequences in the absence of their cognate coding regions (called isolated 3’UTRs or i3’UTRs) opens up considerable avenues for gene regulation not previously envisioned. Each isolated 3’UTR may still bind and interact with micro RNAs, RNA binding proteins as well as other nucleic acid sequences, all in the absence or low levels of cognate protein production. Here we document the expression, localization and regulation of i3’UTRs both within particular biological systems as well as across the transcriptome. As this is an entirely new area of experimental investigation these early studies are seminal to this burgeoning field.

## Introduction

In contrast to the canonical view that an mRNA molecule is comprised of a 5’UTR, a 3’UTR and a coding region (CDS), we and others have found widespread stable expression of mRNA 3’UTRs in the absence of their cognate coding regions (CDS) (1-4) and CDS sequences in apparent absence of their 3’UTRs. These isolated 3’UTRs (i3’UTRs) can be highly expressed, often in reciprocal patterns to their cognate CDS, and are likely to play an important role in gene regulation, as do other long non-coding RNAs (lncRNA) (5-7). With dual in situ hybridization (ISH)-we previously showed non-random differential 3’UTR/CDS expression in many tissues in the embryo and adult. The 3’UTR/CDS ratio for a given gene is not fixed-for example young dopaminergic (DA) progenitors express high levels of *Sox11* CDS compared to 3’UTR but switch and then maintain high *Sox11* 3’UTR levels (2). Cells, then appear to expend high energy to maintain robust, non-random i3’UTR expression which suggests that either the ratio of the 3’UTR to its cognate CDS (3’UTR/CDS), and/or the isolated 3’UTRs themselves may have important gene regulatory functions.

Historically there are some reports of biological roles for 3’UTRs independent of their cognate CDS; found primarily when i3’UTR’s were intended as a control, but where their overexpression or deletion produced a biological effect (8-10). More recently however are a few reports specifically focused on elucidating a particular role for a 3’UTR finding specific functions in axon growth and viability (11). While most i3’UTRs are likely to have a biological role, as with long non-coding RNAs (lncRNAs), these actions/functions may be diverse and cell type contextual (12-16), mediated by binding to RNA, RNA binding proteins, and/or DNA and may be involved in functions ranging from the regulation of stem cell pluripotency to cancer progression (13, 17, 18). For this reason, a thorough understanding of when, where and which mRNAs show high tendency to show differential 3’UTR/CDS expression across tissues and genes sets is useful and necessary to devise cogent hypotheses that can be developed and tested for any individual i3’UTR. Here we present *in vitro, in vivo* data on the non-random and dynamic i3’UTR expression of select genes in progenitor cells, followed by *in silico*, unbiased transcriptome-wide analyses of open source RNAseq genes sets. Togethers these data show a strong proclivity for high 3’UTR/CDS in cells and for genes involved in transition states, during both proliferative and developmental maturation events.

## Results

### Widespread use of differential 3’UTR/CDS usage and ISH specificity

Figure 1 serves to illustrate both the widespread use of differential 3’UTR/CDS expression across tissues and ages, as well as extensive controls carried out to ensure high fidelity of our dual situ hybridization (ISH) procedure. Examining > 60 cognate 3’UTR/CDS probe pairs across tissues, genes and ages we document probe specificity, pattern replication and congruence with public data (Fig. 1;A-G) and show that the 3’UTR/CDS expression of closely related family members, although overlapping is distinct, revealing biological subtleties previously unseen. First from serial adjacent sections we find that multiple *Sox* family members each show distinct ISH patterns with no cross reactivity across family members; *Sox9* and *Sox11* in the developing head show overlapping but unique expression (Fig. 1A-C; arrows). Further, the *Sox11* signal is absent in *Sox11* KO animals, while *Sox2* and *Sox9* expression remains intact (Supp. Fig. 1A-H; C vs D). In the developing mid-hindbran, previous single probe ISH analyses shows *Sox* family member genes to have fairly ubiquitous, overlapping expression with undiscernible patterns (Supp Fig. 1Q’ vs R; http://developingmouse.brain-map.org/experiment/show/100071526) (19, 20). However, dual 3’UTR/CDS ISH reveals dramatic differential expression for *Sox11, 12* and 4 in this region with unique, distinct, finely tuned expression (Fig. 1J-N and Supp Fig. 1L,M; *Sox11* shows higher CDS/3’UTR in progenitor regions similar CDS/UTR in more mature neurons (Fig. 1K; white arrow), and high 3’UTR/CDS in the most differentiated (dorsal) neurons (Fig. 1K; carrot) with other *Sox* genes showing different patterns (Fig. lL-N; carrots)). Such differential and distinct 3’UTR/CDS expression for each gene in developing neurons (as well as other tissues) is likely to be critical during important developmental windows.

**Figure 1.**
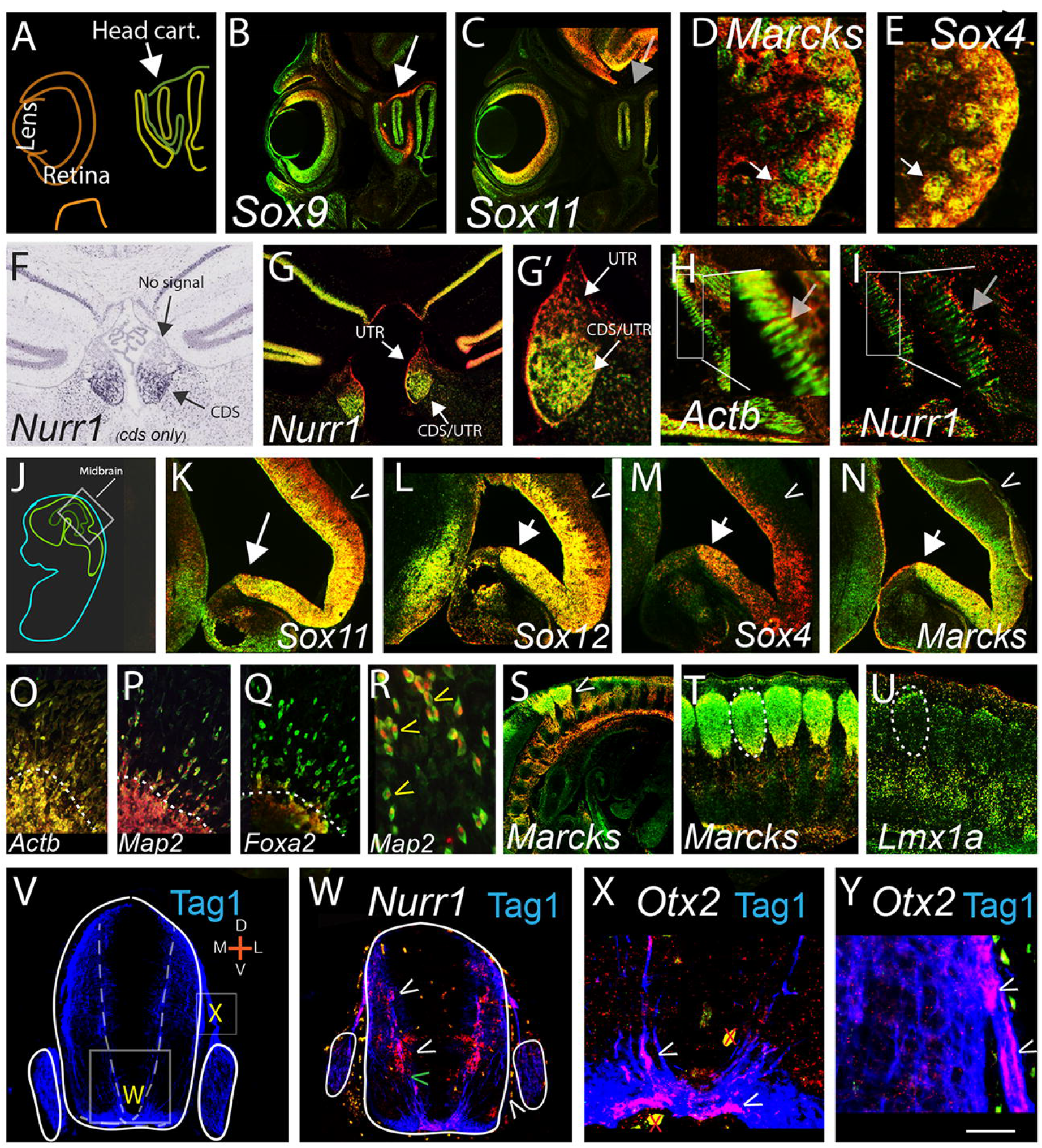
Widespread, specific, differential 3’UTR (red)/cognate CDS (green) expression across mouse tissues and ages. (A) Schematic of sagittal E15 eye and head cartilage region. (B,C) similar sections to A; probes for *Sox9* and *Sox11* detect unique signals (white arrow, signal, gray; no signal) (**unless specified, in all figures 3’UTR probe is red and CDS probe is green with both probes co-hybridized to same section**). (D,E and Supp. Fig.1I,J) Kidney; *Marcks* shows high CDS in glomeruli, while *Sox4* shows equal 3’UTR/CDS in same region in adjacent section; white arrows. (F,G,G’); *Nurr1* in adult brain hybridized with Allen brain atlas single CDS probe (F) (https://developingmouse.brain-map.org/experiment/show/732) or in-house probes (G,G’). (G’) high power view of medial habenula showing high *Nurr1* 3’UTR throughout, CDS restricted ventrally. (I) *Nurr1* in E15 muscle shows distinctly high 3’UTR at muscle fiber tips while *<u>Actb</u>* (H) shows CDS with some 3’UTR throughout; (H,I; gray arrows; insets). (J) Schematic, sagittal E15 embryo. (K-N) Similar adjacent sections as J inset, showing *SoxC* family members and *Marcks*. Arrows show newly developing neurons, carrots show dorsal brain (additional genes in Supp. Fig.1 L,M). (O-Q) *Actb,Map2,Foxa2* 3’UTR/CDS in E15 neural explants *in vitro;* dotted line shows explant edge, cells above that are migrating on substrate. (R) High power showing *Map2* 3’UTR confined to the nucleus with the CDS in the cytoplasm in migrating cells. *Actb* also shows high 3’UTR nuclear signal in migrating neurons, while *Foxa2* (Q) does not. (n=16 explants/probe; all probes showed replicate patterns). (S-Y) 3’UTR/CDS expression in and around developing axons; (S,T) In sagittal view *Marcks* shows high CDS in DRG cell bodies (dotted circle T) but high 3’UTR in axons (carrot; S). (U) *Lmx1a* in adjacent section shows no obvious axonal expression. (V-Y), E11 spinal cord-early developmental genes show high 3’UTR in or alongside Tag1+ axons (V). (W) *Nurr1* 3’UTR is highly expressed along axon path; carrots. (X,Y) *Otx2* in the axons; carrots. Both *Otx2* and *Nurr1* show robust CDS signal in other sections (G, and Supp. Fig.1O or not shown). Scale bar, A-C; 600, D,E; 300 F,G; 450, G’ 200, H,I; 1200; inset 300 J; 1600 K-N; 300, O-Q 800, R; 300, S; 1500, T,U; 500, V,W; 220, X, 100, Y, 80 um. Red x’s in X are artifacts.

We further compared in-house generated probes to custom RNAscope^r^ probes and found indistinguishable hybridization patterns between the two (not shown) and verified that co-hybridization with identical probes labeled with the two different fluorphores-digoxygenin or fluorescein-resulted in the majority of positive cells appearing yellow, as expected from signal overlap (not shown). We did find slight non-overlap at tissues edges, thus all data herein is from non-edge tissue regions. For downstream analysis we routinely confirm that ISH signals replicate bilaterally, across sections and in multiple experiments and that dissimilar genes in serial sections show unique hybridization patterns as shown in the developing kidney (Fig. 1D-E; arrows and Supp. Fig. 1I,J). Our dual ISH procedure is therefore specific and sensitive and can distinguish differential signal in closely juxtaposed cells (Fig. 1). Moreover, comparing our data to the widely used Allen Brain Atlas we find identical expression between the two for *Nurr1* (as well as other genes in the atlas). Both Allen brain (Fig.1F) and in-house (Fig. 1G,G’) CDS probes detect signal in the ventral, but not dorsal, medial habenula. Additional analysis of *Nurr1* expression shows that in contrast to the restricted *Nurr1* CDS, the *Nurr1* 3’UTR shows widespread expression including in developing muscle, spinal cord and adult cerebral cortex, (Fig. 1I and Supp. Fig. 1K,S). In the CNS *Nurr1* is known primarily for its role in dopaminergic (DA) neuron development but these data suggests a wider role in the nervous system, as well as a role in other tissues (see Discussion).

These data demonstrate that our ISH procedure to specific and sensitive, and reveal an extremely penetrant use of differential cognate 3’UTR/CDS expression in developing and adult, neural and non-neural tissue for many/all genes examined.

### Cellular compartmentalization of the i3’UTRs

As shown *in vivo* previously (1, 2), when both the CDS and cognate 3’UTR are expressed in a cell, they may be localized to different compartments (ie nuclear vs. cytoplasmic). We asked here if this localization is fixed or dynamic, comparing aggregated to migrating embryonic peripheral neurons. Both *Map2* and *Actb* (each are structural proteins) show nuclear localization of their 3’UTR, for *Actb* this occurs in both aggregated and migrating neurons, but for *Map2*, there is a switch from 3’UTR/CDS nuclear and cytoplasmic overlap in aggregated neurons, to sequestration of the 3’UTR to the nucleus and CDS to the cytoplasm in migrating neurons (Fig. 1O-R; R carrots). Thus, localization of 3’UTR and CDS sequences is dynamic, varies from gene to gene and is dependent on the cellular state. Of more than sixty genes examined, most showed differential 3’UTR/CDS across cellular compartments in vivo or in vivo, in at least one tissue.

Neurons are unique due to their extended axonal processes, and many important cellular processes take place in the axons including translation of mRNA (21-23). Of more than 20 genes examined in axons four showed prominent differential 3’UTR/CDS in or around axons (Fig. 1S-Y), *Nurr1* shows extreme high i3’UTR expression in cells just lining spinal cord TAG1+ axons (Fig. 1W), while *Otx2* (Fig. 1X,Y) and *Wnt1* (not shown) show high i3’UTR expression within these axons, in the absence or with low levels of CDS (Supp. Fig.1 N-P; see O). In contrast the *Marcks* gene shows high 3’UTR expression in peripheral dorsal root ganglion neurons, while the cell bodies of these neurons produce predominantly *Marcks* CDS (Fig. 1S,T). Thus i3’UTRs may be specifically localized to cell bodies and/or axons.

### Differential 3’UTR/CDS in proliferating cells

Due to the strong use of differential 3’UTR/CDS expression in genes involved in developmental processes (2), we examined the 3’UTR/CDS expression of the critical early pluripotent genes (PpGs); *Myc, Nanog, Sox2, Oct4*, and *Klf4*. Each PpG was compared to 11 other genes in two systems, the E12 olfactory epithelium (OlfE) (Fig. 2A), as well as the developing whisker and hair niche (Fig. 2H and 3). In the OlfE subventricular zone (SVZ) all PpGs are expressed and most show striking differential 3’UTR/CDS expression (Fig. 2B-G and Supp. Fig. 2A-H). In low power *Sox2* and *Ascl1* (a pro-neural gene used as a control) show unique 3’UTR/CDS patterns across the OlfE; *Sox2* with graded differential 3’UTR/CDS; and *Ascl1* with more coincident 3UTR/CDS expression-both show identical patterns bilaterally (Fig. 2B,C;*). Within the SVZ neural progenitors are located apically, more differentiated cells basally, and radial glia-labeled by Nestin-span the distance. We asked, then if PpG 3’UTR/CDS expression ratios correlate with these subregions. All the PgGs except *Myc* show preferential localization of their 3’UTR to apical cells, with isolated CDS throughout the radial glia (labeled by Nestin) into the basal regions (Fig. 2E-G and Supp. Fig. 2G,H). For *Myc* both the 3’UTR and CDS are largely restricted to the SVZ apical progenitor region (Fig. 2E; yellow carrot, Supp. Fig. 2E).

**Figure 2.**
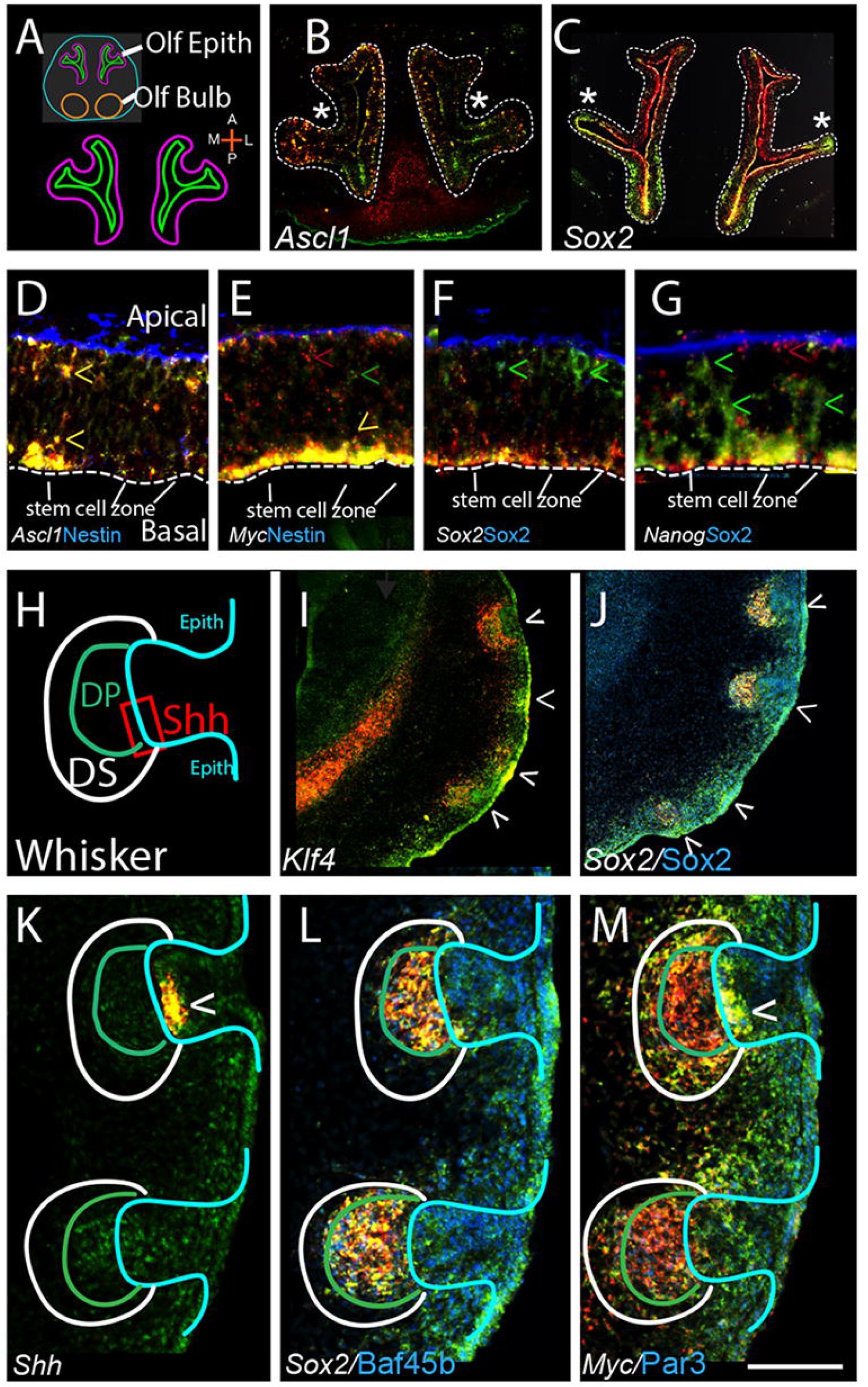
Progenitor cells show distinctive 3’UTR/CDS patterns. (A) Schematic showing horizontal view of E12 olfactory epithelium (OlfE). (B,C) similar sections of E12 OlfE with control pro-neural gene *Ascl1* and PpG gene *Sox2. Ascl1* is expressed in scattered cells, often with colocalized 3’UTR/CDS expression, while *Sox2* shows graded differential 3’UTR/CDS expression across OlfE regions. * denotes symmetrical patterns in contralateral OlfE for each gene, (additional genes in Supp. 2 Fig. B,D-H). (D-G) higher power view of core PpG expression in OlfE SVZ; as labeled. Note that all PpGs (but not *Ascl1* (D)) show preferential localization of their 3’UTR sequences to the apical plate, labeled in D and same in E-G. Basal plate shows high Nestin (D,E) and Sox2 (F,G) protein and each CDS can also be colocalized to Nestin positive radial glia (seen at longer exposure) (green carrots). Apical stem cell zone is marked. For *Myc* both the 3’UTR and CDS are concentrated apically. For all other PPGs examined, *Sox2, Oct4, Klf4*, and *Nanog*, their 3’UTR expression is concentrated apically with the CDS signal both apical and extending along radial glia to the basal plate; carrots; F,G and Supp. Fig. 2). (H) Schematic of whisker key anatomy. (I,J) low power views of four developing whiskers; top two with well-defined dermal papillae, bottom two-earlier condensates. Note that differential 3’UTR/CDS expression for all PpGs (I, J and Supp. Fig. 2M,N) is apparent and an early indicator of hair peg condensation. (K-M) higher power views showing differential 3’UTR/CDS for *Sox2* (L) and *Myc* (M) in anatomically distinct regions of the whisker. *Shh* (K) shows coincidental 3’UTR/CDS expression in an adjacent section (carrot). Other PgPs shown in Supp. Fig. 2I-L). Anatomical abbreviations are, Shh; *Shh* expression region, DP; dermal papilla, DS; dermal sheath (white). Scale bar, A; 500, B,C; 350, D-G; 150, H,K-M; 55, I,J; 350 um.

Because whisker (Fig. 2H) and hair follicle niches (Fig. 3A-C) are stereotyped structures where progenitor, proliferative, and other cell domains have been defined anatomically (24, 25) we asked if 3’UTR/CDS expression correlates with anatomical structure and/or cell proliferation. At postnatal day 2, differential PpG 3’UTR/CDS expression is detected in whisker sub-regions even in early condensates (Fig. 2I,J; carrots show developing whiskers). In the more mature whiskers with clearly visible dermal papilla (DP) there is differential 3’UTR/CDS expression for all the PpGs (Fig. 2I,J; top two carrots and Supp. Fig. 2M,N). At higher power, it is clear that *Myc* shows equal 3’UTR/CDS in the proliferative epithelial region (the Shh+ cell region (Gonzales and Fuchs 2017) and *Shh* 3’UTR and CDS positive in Fig 2K,M; carrots). In contrast, in the closely apposed but distinct dermal papilla, *Myc* shows very high levels of 3’UTR vs. CDS expression (Fig. 2M; inside green circle) while *Sox2*, a gene necessary for DP differentiation (26) shows equal 3’UTR/CDS (Fig. 2L; other PpGs in Supp. Fig. 2I-L). The juxtaposition of dramatically different 3’UTR/CDS ratios for the PpGs in closely apposed anatomical structures emphasizes that 3’UTR/CDS ratios are very tightly regulated, showing different or reciprocal relationships in distinct biological structures. Importantly, the 3’UTR/CDS ratio for the PpGs may be critical in developmental events such as neurogenesis and hair formation.

**Fig. 3.**
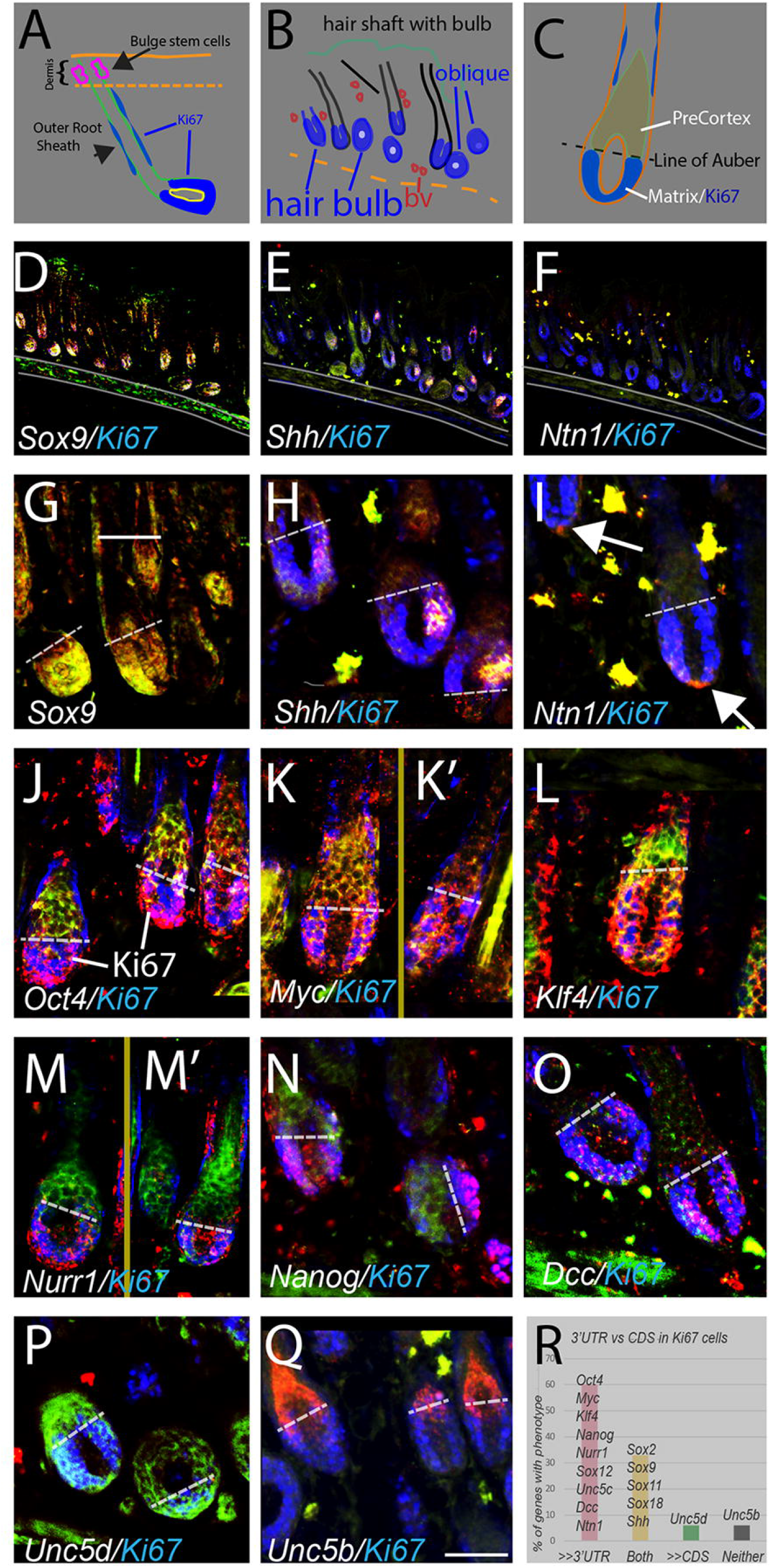
The 3’UTR/CDS ratio for PpG and other genes changes abruptly at the boundary between the Ki67+ proliferative zone and non-proliferative zone. (A) Key anatomy of skin hair follicle; stem cells located dorsally within dermis, outer root sheet (ORS) at the perimeter of the follicle, proliferating cells (Ki67+) in deep hair bulb and along ORS. (B) schematic of skin sections shown in (D-F), each section shows multiple hair follicles; oblique deep bulb follicles appear round. (C) coronal view of lower hair shaft showing location of Ki67+ matrix cells, and pre-cortex cell. The critical line of Auber (dotted line in C-Q) demarcates the boundary between proliferating Ki67+ cells and non-proliferating pre-cortex cells. (D-F) similar sections as B hybridized for *Sox9, Shh* and *Netrin-1 (Ntn1*). Note the different (and therefore specific) hybridization patterns for the different probes. (G-Q) higher power views of 1-3 hair bulbs for genes as labeled. Note abrupt shift from high 3’UTR in Ki67+ cells below line of Auber to Hi CDS above. (R) 60 percent of genes examined show high i3’UTR in Ki67+ cells with a switch at the line of Auber (n=20-30 bulbs/gene). (K,K’, M, M”) cropped to remove hair shafts which can show high autofluorescence. Full picture of M can be seen in Supp. Fig. 3H. Second bulb in N is oblique (as in A) therefore line of Auber appears rotated. Scale bar, A; 160, B; 400, C; 220, D-F; 700, G-Q; 85 um.

To closely examine the relationship between differential 3’UTR/CDS ratio and cell proliferation we examined deep hair follicles-above and below the “critical line of Auber”; cells below the line are proliferative Ki67+ cells, above are the non-proliferating “pre-cortex” cells. Based on the strong localization of PpG 3’UTR expression to neural stem cells we hypothesized that PpG’s may show high 3’UTR below the line of Auber. First, across serial adjacent tissue sections we find very different patterns of expression for the individual genes and each pattern replicates across hair follicles within the section (Fig. 3D-F and Supp. Fig. 3A-C). Secondly, we found dramatic and abrupt shifts in 3’UTR/CDS expression at the line of Auber for *Oct4, Klf4* and *Nanog* (as well as *Nurr1*and *DCC*) shifting from high i3’UTR expression in Ki67+ cells, to high CDS in pre-cortex cells above the line of Auber (Fig. 3J,L,N and M; dotted line is line of Auber). *Myc* showed more coincident expression but also higher 3’UTR than CDS in Ki67+ cells compared to pre-cortex (Fig. 3K,K’). In contrast *Sox9* and three other *Sox* genes showed similar 3’UTR/CDS throughout (Fig. 3G and Supp. Fig. 3F,G), while *Shh* showed asymmetric localization, and *Ntn1* to the ventral midline, both in subsets of the Ki67+ cells (Fig. 3H,I and Supp. Fig. 3D,E). We also examined the Netrin-1 receptors *Dcc*, and *Unc5B-D*. Like the PpGs, *DCC* (also known as a tumor suppressor gene) showed higher 3’UTR/CDS in Ki67+ cells (Fig. 3O). In contrast to PpGs, the *Unc5c* and *d* receptors showed either no expression, or high CDS in Ki67+, with high CDS or high 3’UTR above the line, respectively (Fig. 3P,Q; and see R). In the nearby blood vessels both *Ntn1* and *Unc5b* show high CDS and high 3’UTR as expected (Fig. 3I and Q; yellow signal). Interestingly, and unexpectedly, *Unc5d* shows very high 3’UTR in these bvs as well (Fig. 3P and Supp. Fig. 3C). For each of the 16 genes examined expression was independently confirmed by scRNAseq (27) or by Tabula Muris (28).

### Cell state and 3’UTR/CDS ratio

While dynamic differential 3’UTR/CDS use is highly prominent during development we asked whether dynamic changes in 3’UTR/CDS ratios can be induced by cell-state changes such as the response to stress. For this we examined NIH3T3 (3T3) cells plated on prepatterned glass chips (Cytoo^r^) (Fig. 4A; in low confluence cultures the cells attach only to the substrated circles). In the smallest circles (80 um)-we find dramatically, that cells in the center of each circle show high Actin (*Actb)* CDS/3’UTR while the cells at the perimeter all show high 3’UTR/CDS, with pattern replication across more than 30 circles/chip (Fig. 4B,C). In confluent cultures in which cells grow onto the glass (Fig. 4 D,E) we examined *Actb* 3’UTR/CDS ratio in stressed vs non-stressed cultures. In the absence of stress most 3T3 cells express higher *Actb* 3’UTR/CDS, except interestingly the cells on the perimeter of the circles (Fig. 4D; white carrot). After stress, most cells switch and now express high *Actb* CDS/3’UTR (Fig. 4E). Assayed again, we compared the 3’UTR/CDS ratios for three different genes, *Actb Map2* and *Sox12*, and find that for each of these genes there is an increase in their CDS levels compared to their 3’UTR levels after stress (Fig. 4 F-K).

**Figure 4.**
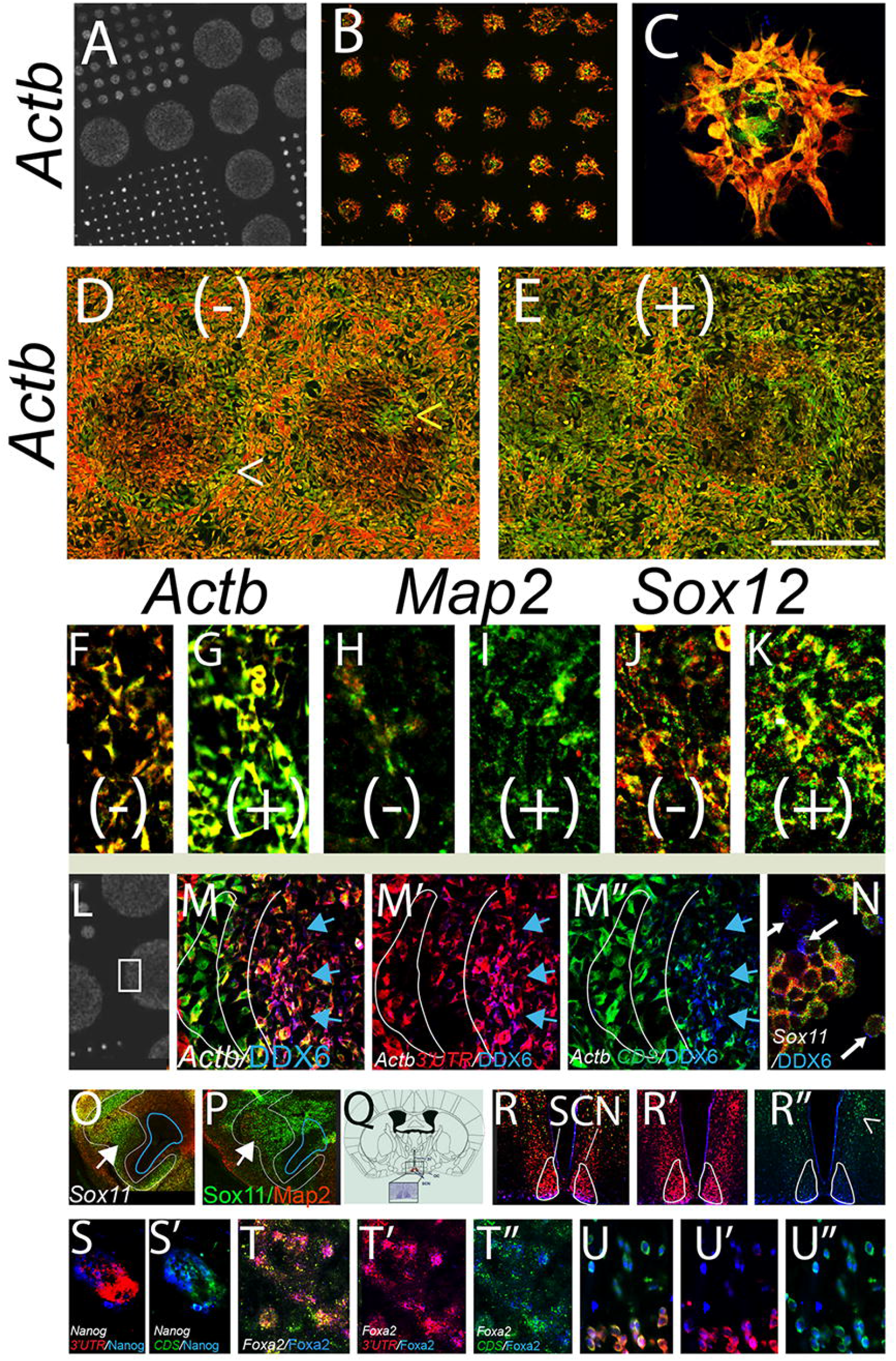
3’UTR/CDS ratios change with cell state. (A) Photo of Cytoo^r^ chip with circles 800 to 80 um arranged across chip. (B,C) In 80 um circles, 3T3 cells consistently show high *Actb* CDS in center cells and high 3’UTR in perimeter cells (technical replicates n >30-50 circles/chip, biological replicates n=2). In highly confluent cultures cells grow outside the circle. Under normal conditions (D) all cells show very high 3’UTR/CDS except cells at circle border that show high CDS/3’UTR. Stress challenge with 30 min sodium azide (NaZ) (E; + denotes NaZ) causes most cells to express higher *Actb* CDS compared to 3’UTR (D,E are matched photos). Mini-islands of cells that express the opposite ratio from the majority can be seen, suggesting that the 3’UTR/CDS ratio is due to the state of the cell, not to a hybridization artifact of the chip (Fig. 4D; yellow carrot). For three different genes, all increase their 3’UTR/CDS ratio after stress, (F,G) *Actb*, (H,I) *Map2*, (J,K) *Sox12* (n=10 technical replicates, 2 biological replicates). (L-N) Localization of P-body phase marker DDX6 in relation to 3’UTR/CDS. (L) region of Cytoo^r^ chip shown in M-M”. DDX6 is only found in cells with *Actb* 3’UTR (M’; *Actb* 3’UTR red, DDX6 blue), not with cells that express primarily CDS (M”; *Actb* CDS green, DDX6 blue); blue arrows. In Neuro2A cells neither the 3’UTR nor CDS of *Sox11* colocalizes with DDX6 (N; white arrows show DDX6 label in blue). (O,P) Sox11 antibody stain shows same pattern as *Sox11* CDS in similar but non-adjacent section (white arrows and outline; blue outline shows ventricle) (outline from O superimposed onto P). (Q) schematic showing location of suprachiasmatic nucleus SCN (clock center) in the adult brain. (R-R”; *clock* 3’UTR red, CDS green, clock protein, blue). Note that Clock protein shows greater co-localization with the *c lock* 3’UTR (in SCN outlined in white), than with cells with high *Clock* CDS; R” carrot. (S,S’) *Nanog* mRNA and protein in hair bulb (S,S’; *Nanog* 3’UTR red, CDS green, nanog protein blue). Note that in these cells nanog is is more highly co-localized with the *Nanog* CDS than 3’UTR. (T-U”) *Foxa2* 3’UTR, red, CDS green, protein blue. (T-T”; embryonic dopamine neurons, U-U”; embryonic dorsal root ganglion neurons). Note that the Foxa2 protein localizes with the *Foxa2* 3’UTR in DA neurons but equally with 3’UTR and CDS in migrating DRG neurons. Scale bar, A; 1700, B; 450, C; 90, D-E; 600, F-K; 150, L; 1200, M,M’, M”; 300, N; 180, O,P; 650 Q; 5700, R, R’, R”; 900 S, S’75, T,T’,T” 3000, U; 430 um.

### i3’UTR or CDS co-localization with other markers

Of interest of course is how 3’UTR sequences (or CDS sequences) become sequestered within cells. For this we examined the co-localization of various probe sets to specific antibody markers for the nuclear membrane (Lamin), P-body phase regions (DDX2, Rcd2, xrn1), endosomes (Lamp1), and dead cells marked by activated Caspase3. Unlike Ki67 which is highly correlated with the i3’UTR for many genes, we saw no preferential localization of 3’UTR or CDS for activated Caspase3 or Lamp1 for any gene examined (2-4 for each protein). Because mRNA is often localized to P-bodies we examined DDX6 and other Pbody markers. We found that in 3T3 cells, the *Actb* 3’UTR (with or without CDS) was highly colocalized with P body markers (Fig. 4L-M”; shown is DDX6”), while the CDS only showed co-localization in the presence of the cognate 3’UTR (Fig. 4M”). While striking for *Actb* in 3T3 cells, neither the *Sox11* 3’UTR nor CDS showed co-localization with DDX6 in 3T3 or Neuro2A cells (Fig. 4N; shown is Neuro2A). Thus, while 3’UTR sequences may co-localize with P bodies (with or without the CDS), this is not a requisite location. We saw similar but less robust colocalization of some 3’UTR sequences with Lamin. Because these localizations appear gene and cell dependent, a more extensive analysis was not pursued here, however analyses in individual biological contexts is likely to be informative.

Equally important is how 3’UTR/CDS ratios correlate with cognate protein levels. We reported previously that for two cytoplasmic proteins, a high 3UTR/CDS ratio correlated with low protein expression (2). Upon further investigation we find that in the systems outlined here Sox2, Sox11 and Nanog proteins are also often likely detected in cells with high or equal CDS/3’UTR vs high 3’UTR/CDS (Fig. 4O,P, S,S’ and Fig. 2F). However, after examining multiple probe sets and cognate proteins; in some cases, especially for transcription factors we found preferential localization of the protein with the cognate 3’UTR, this was true for Clock and Foxa2 (Fig. 4R-R”, T-U”), Klf4 and Oct4 (not shown), as well as others. Foxa2 protein is highly co-localized with the *Foxa2* 3’UTR in embryonic DA neurons in culture but not in same age peripheral neurons (Fig. 4T-T” vs U-U” respectively). While unexpected that some proteins would show higher colocalization with cells that are expressing higher cognate 3’UTR (than CDS), it was observed across multiple experiments and cell types with a number of different antibodies to different proteins. (This was not an antibody artifact as not all antibodies showed such co-localization with the 3’UTR sequences). Because this is highly surprising and differs across proteins, and cell types, it is beyond the scope of these studies to describe protein levels in every context mentioned in this report; each will require specific, detailed study, and generalizations would only hamper our forward knowledge. However, it is striking that in many cases, protein is more highly localized to cognate hi 3’UTR compared to hi CDS cells.

### In silico; i3’UTRs across tissues

The ISH studies here and by others strongly suggest that one gene utilizes different 3’UTR/CDS ratios across tissues and that across development, and that any particular gene may change this ratio over time. We turned then to verify these findings in an unbiased manner examining open source RNAseq data. In all bulk (Figs.5 and 6), and single cell data sets (not shown) examined (> 50 data sets) we find this to be the case. For example, comparing multiple *Sox* gene family members across 11 randomly chosen data sets we find higher 3’UTR/CDS expression in differentiated tissues (Fig. 5A; teal bar), but a striking reversal in embryonic stem cells (ESC) (Fig. 5A; pink bars) in which there is higher CDS/3’UTR expression. Conversely, to examine changes in ratio of 3’UTR/CDS for a given gene in one tissue over time, we queried different aged heart tissue (Fig. 5B) and found, for example that *Hnmpl* shows high 3’UTR/CDS in the embryo but low 3’UTR/CDS in the adult (Fig. 5B; top traces). In contrast, *Dusp3* shifts from high 3’UTR/CDS in the embryo to comparable levels in the adult (Fig. 5B; middle traces). Thus, as predicted from ISH studies genes change their ratio depending on the cell state, and/or maturation level. All heart data sets were generated by the same investigator, ruling out the possibility that differences are due to library generation or sequencing methodologies.

**Figure 5.**
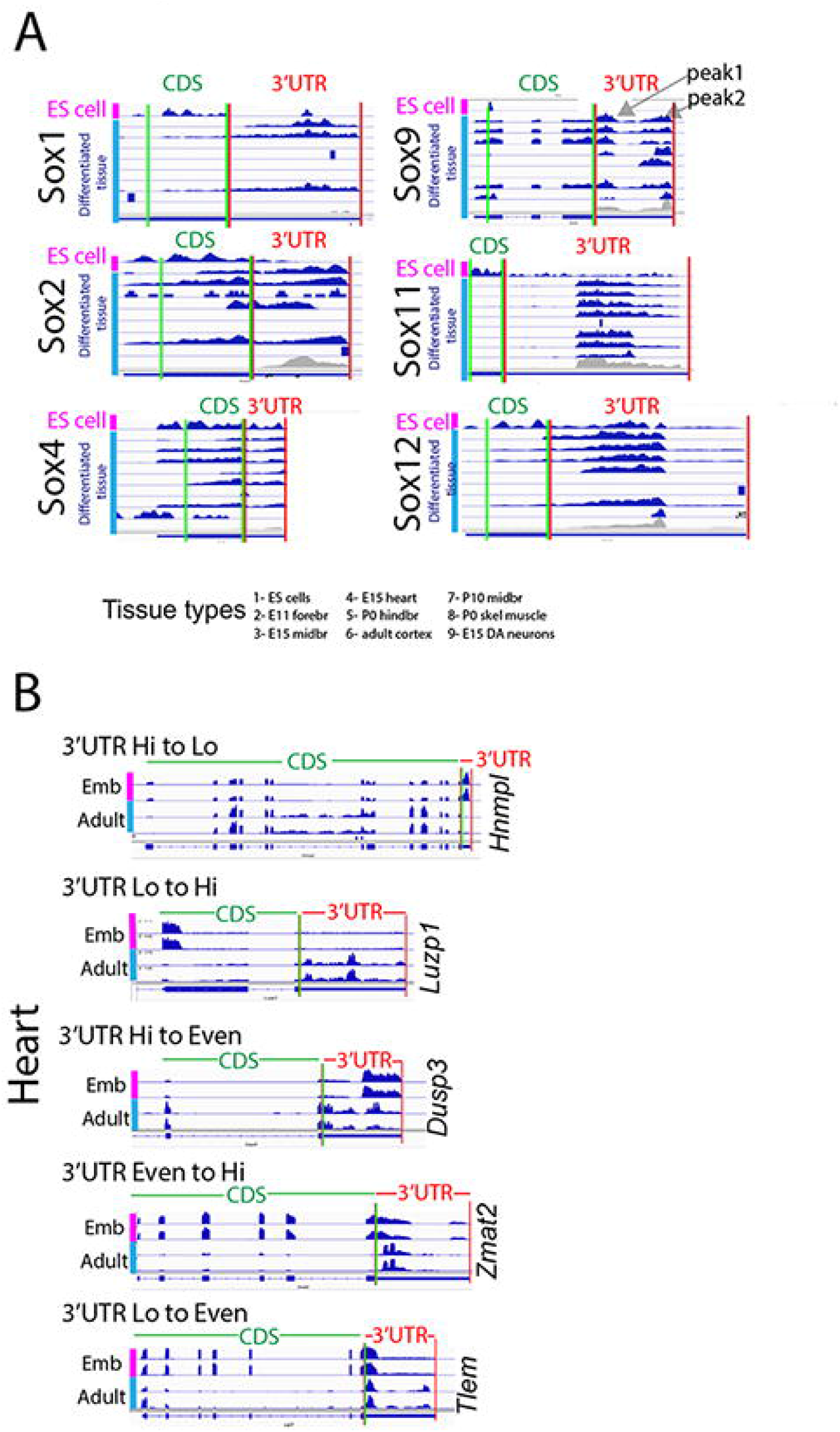
Widespread differential 3’UTR/CDS use is reflected in RNAseq data. (A) RNAseq data from 9 tissues showing 3’UTR and CDS expression for select *Sox* genes. For each gene, red vertical lines denote 3’UTR region, green denote CDS. Top trace for each gene (pink sidebar) is embryonic stem cell (ES) cell. Next eight traces (blue sidebar) show differentiated tissues, the first seven from open source Geo; noted at bottom, and the last, in-house RNAseq from E15 dopamine neurons (2). As noted in (1, 2) and shown in gray, RNAseq traces are not uniform across the entire 3’UTR and can be divided into “peaks” and “valleys” (eg see *Sox9*; peak1 and peak 2; valley in between) (see text for definition). Each *Sox* gene shows greater (or in one case equal) 3’UTR to CDS expression in differentiated tissues. ES cells show the opposite, with high CDS/3’UTR. (B) Representative genes from embryonic (Emb; pink bars) and adult (Adult; blue bars) heart databases (each with two replicates) illustrate 3’UTR/CDS ratio changes over time; e.g., *Hnmpl* 3’UTR/CDS changes from high to low, *Luzp1* from low to high. CDS - 3’UTR junction marked by double red-green line and labels above.

### Classes of genes that show high 3’UTR/CDS

We then asked in a general sense what particular genes sets or categories (GO categories) tend to show a high, or low 3’UTR/CDS ratio. We first examined two tissues; hindbrain and heart across multiple ages. Each sample had two biological replicates (with high reproducibility across replicates in genes “called” Hi 3’UTR; 63% to 88% identity) and we asked, for each tissue and age, whether the GO categories corresponding to Hi (>0.6) or Lo 3’UTR genes (< 0.4) (see methods for details) are different. We found that first, regardless of tissue or age the top GO category for Hi CDS (Lo 3’UTR) genes is consistently “translation”, followed by “ribonucleoprotein complex biogenesis” (Fig. 6A; left). On the other hand, intriguingly, the GO categories for Hi 3’UTR genes *are remarkably specific to both the age and function of the target tissue*. In hindbrain, the high ranking GO categories are “embryonic morphogenesis”, “neuronal differentiation”, and “synaptogenesis” E15 to P0 tissue. Likewise, but heart specific, the top high 3’UTR GO categories for heart are “actin filament based processes” “heart development, “actin filament”, and “Gtpase signal transduction” from E15 to P56 (Fig. 6A; right columns). The GO categories of Hi 3’UTR genes then are representative of cell and tissue type with high representation of cell fate determination genes. This, regardless of the fact that both Hi and Lo 3’UTR genes each represent less than 10% of all the genes expressed in that tissue and are not necessarily the most highly expressed genes.

**Figure 6.**
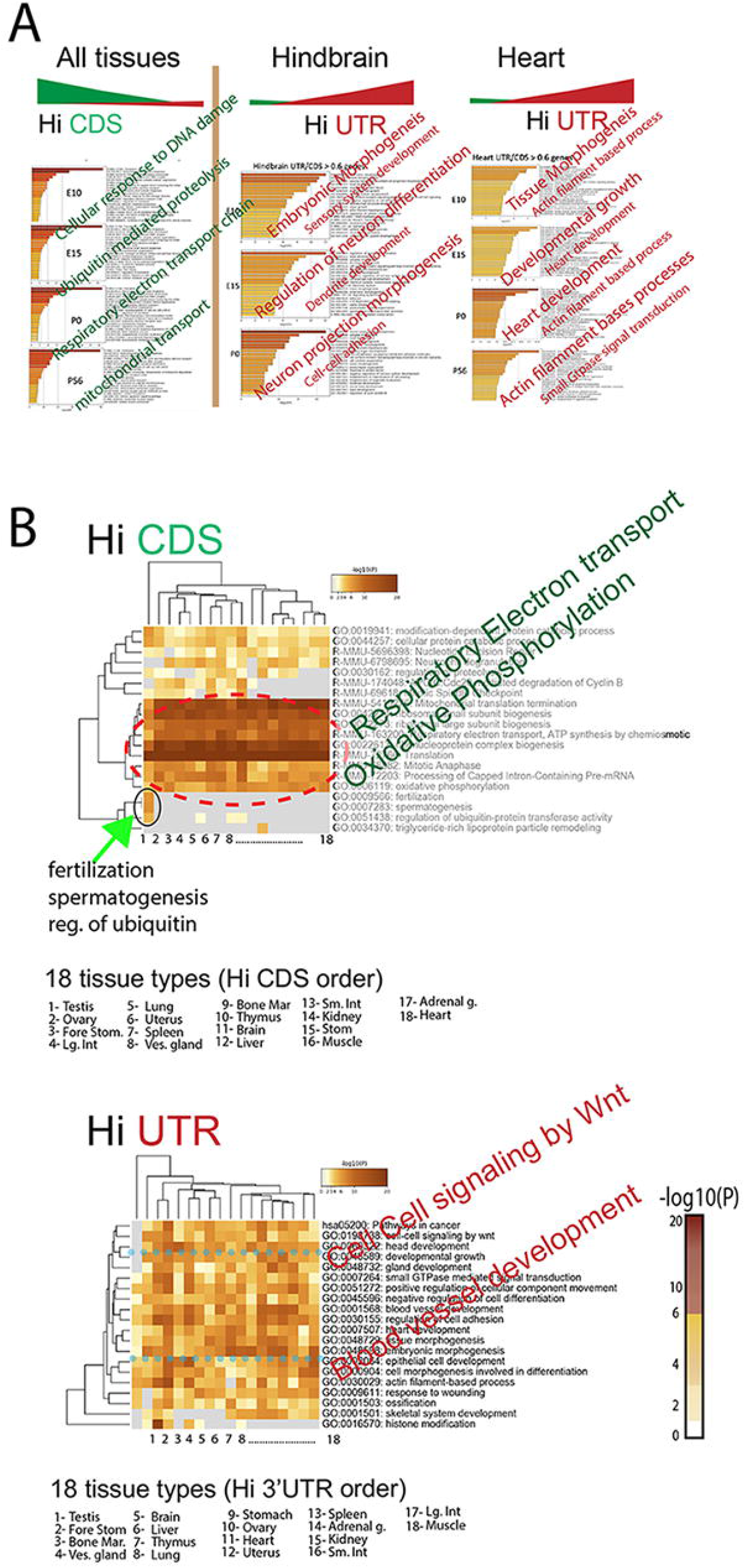
Unbiased analysis shows that Hi 3’UTR/CDS genes define tissue specific properties. (A) Gene Ontology (GO) categories for Hi CDS and Hi 3’UTR (see methods for gene selection criteria) genes for hindbrain and heart data sets. Right columns; top GO for Hi 3’UTR genes, left column; top GO for Hi CDS genes. GO categories are listed in black with examples shown in larger colored diagonal text. (B) Top GO categories across 18 tissues-heat map shows significance for each tissue. Order of tissues for Hi CDS is different from Hi UTR; see legend. In all cases the top Hi CDS GO (dotted red circle) categories are oxidative phosphorylation, electron transport and other basic cellular mechanisms. Hi 3’UTR GO categories vary by tissue, are tissue specific and show no obvious pattern (bottom heat map); blue dotted line shows significance variability across tissues for two representative GO categories. Of note, testis Hi CDS GO categories include fertilization and spermatogenesis in addition to basic cellular mechanisms; green arrow, top heat map. Further, testis Hi UTR GO categories are less aligned with other tissues; bottom heat map, column one, note the gray squares.

While intriguing, we wanted to see if this was wide-ranging and therefore examined this across 18 tissues-with similar results-across all each of the 18 tissues the top GO categories for Hi CDS genes were “respiratory electron transport” and “oxidative phosphorylation” (Fig. 6B; top). And again, we found that high 3’UTR GO categories differ by tissue, with few commonalities between them (Fig. 6B; bottom heat map), and with GO categories highly representative of tissue type. For example the GO category “epithelial cell development” is significant in liver, with “blood vessel” in small intestine and kidney (Fig. 6B). While we consistently find that the high CDS genes are similar in all tissues and that the top categories are primarily involved in cellular maintenance we also find that some categories, although not the most significant, are significant and tissue-specific. In testis for example two significant Hi CDS categories are “fertilization and spermatogenesis” (Fig. 6B; green arrow; black circle). Liver also shows significance for “triglyceride-rich lipoprotein particle remodeling”. Thus, apart from a preponderance of Hi CDS use amongst cell maintenance genes, there is also some tissue specific use of Hi CDS. Testis is an outlier for high UTR GO categories as well, and does not show any of the same GO categories as other tissues (Fig. 7C; bottom; black circle).

**Figure 7.**
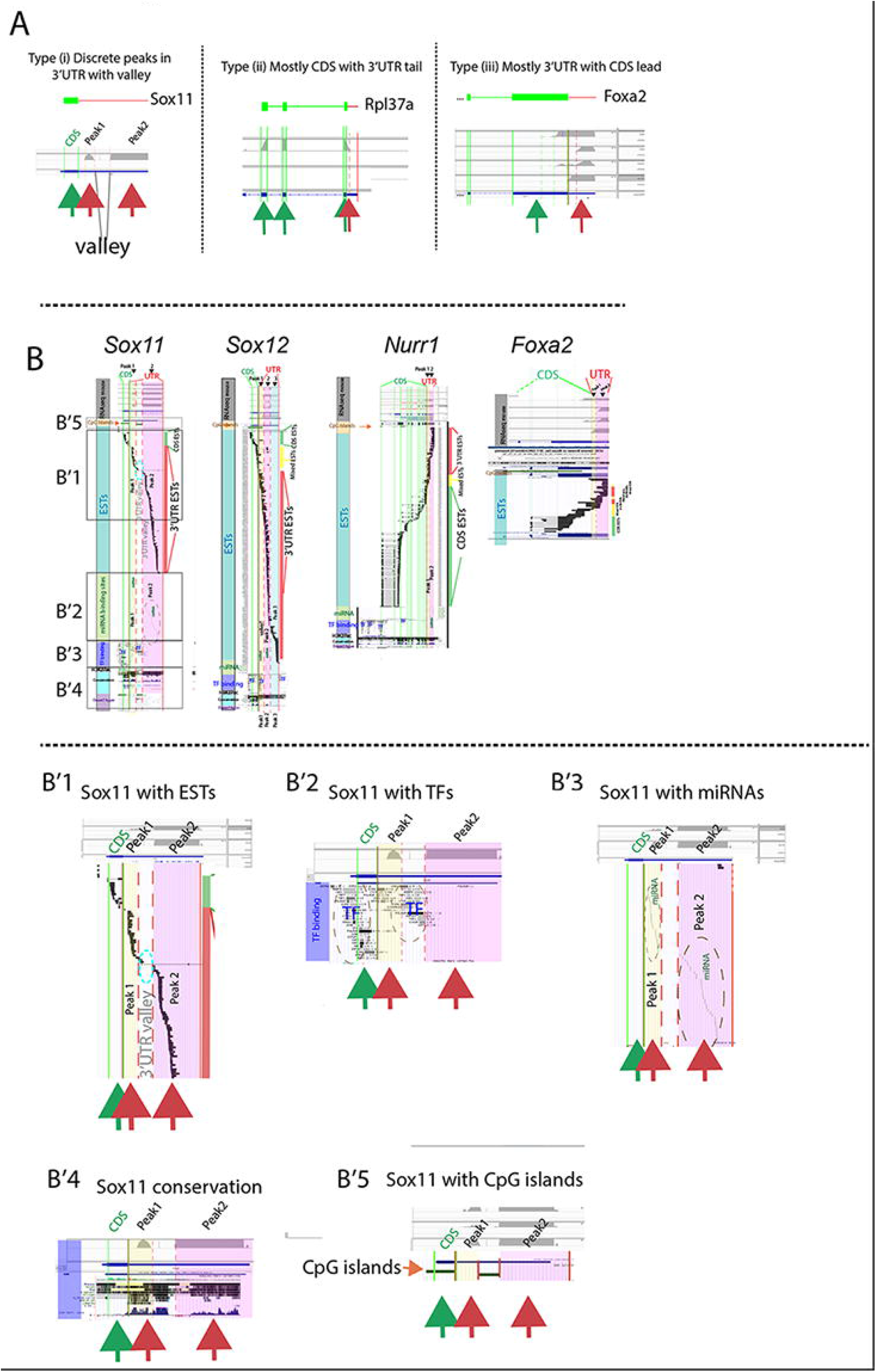
Gene elements and EST sequences correlate with 3’UTR sequence peaks and valleys. (A) illustrative examples of 3’UTR vs CDS peak and valley expression. Gene structure at top. Mouse E15 DA neuron RNAseq trace below. Solid vertical green/red lines denote CDS or 3’UTR; dotted lines denote ends of RNAseq traces. (B) For 4 representative mouse genes; CDS, 3’UTR and prominent RNAseq 3’UTR peaks and valleys are aligned with human gene elements from the UCSC browser **http://genome.ucsc.edu/**; human Hg19. Human data was used since it contains more complete gene element data. For each gene, color-coded vertical bar denotes elements. Top, gray; E15 midbrain mouse RNAseq traces (2), Orange; CpG islands, Teal; expressed sequence tags (ESTs), Green; miRNA binding sites, Purple; transcription factor (TF) binding sites, Brown; H3K27ac binding sites, Light blue; conservation across species, Lavender; Dnase1 hypersensitive sites (Dnase1 hyper). *Sox11* is first gene on left. Below, (B’1-5) are expanded views of *Sox11* data from regions in boxes. Peaks are defined as continuous regions of >150 bp of sequence with FPKM>100 in 4 replicate data sets, valleys show the opposite. Peaks and valley denoted throughout. Peak1 is overlain in yellow, peak2; pink, peak3; blue, valleys are white. Dark red, green and yellow vertical bars to the right of each gene denote ESTs that are uniquely 3’UTR (red, no CDS), uniquely CDS (green, no 3’UTR) or mixed (ESTs that span junction, yellow). Note that *Sox11* has no mixed 3’UTR ESTs, and of 122 total ESTs only 10% have any CDS sequence. No ESTs span the 3’UTR valley; blue dotted circle (B’1), TF binding sites are situated between peaks (B’2), miRNA binding is exclusively within peaks (B’3), conservation is high in peak, and valley regions (B’4), and H3k27ac and CpG islands are in valleys or CDS region not in the peak regions (B and B’5). *Sox12* shows similar profile to *Sox11* except that Peak1 is not as well defined and shows mixed peak/valley characteristics. *Nurr1* shows many more CDS ESTs than 3’UTR ESTs with miRNAs and high conservation only in peak2. Dotted red lines denote 3’UTR peaks, dotted green lines, CDS peaks. *Foxa2;* aligned with ESTs only, is shown to illustrate that not all genes resemble *Sox11* and *12*. For *Foxa2*, CDS ESTs outnumber 3’UTR ESTs with many mixed. Human data was used as mouse data for various elements was unavailable.

### i3’UTR and gene sequence elements

Generation of isolated 3’UTR transcripts may occur by de novo transcription or post-transcriptional processing of mRNA, and indeed it may differ for different genes. Here we examine, and find high correlation of specific gene elements with 3’UTR “peaks” for select genes. As seen in Fig. 5A (for example *Sox9*; gray arrows) and shown in (1, 2), RNAseq expression within a given 3’UTR region is not necessarily contiguous with the CDS, nor continuous along the entire annotated 3’UTR but instead there are peaks of expression (defined in Fig. 7 legend). From visual analysis we observe three main patterns across genes, with any one gene showing high pattern reproducibility across tissues (Fig. 7A). Type (i) shows multiple discrete 3’UTR peaks that originate and terminate in the 3’UTR proper, with valleys (absence of peaks) between peaks, Type (ii), sequence immediately distal to the end of the CDS spanning continuously into a short 3’UTR and Type (iii), a long 3’UTR with a short stretch of leading CDS. Because in many cases one gene will show very similar patterns of 3’UTR peaks and valleys across tissues and data sets (Fig. 5 and 6A; see *Sox11*) we examined the possibility that there are gene sequence elements upstream to, or aligned with these peaks and valleys. Using data collected from the UCSC browser (http://genome.ucsc.edu/, human data was used) we aligned conservation across species, transcription factor (TF) binding sites, CpG islands, and miRNA binding sites to 3’UTR RNAseq peaks (mouse data). Remarkably, for some genes, all of these elements show tight correspondence with discrete 3’UTR peaks. For example, for *Sox11* (selected since the location of the 3’UTR peaks and valleys are highly consistent across tissues), CpG islands, H3K27ac peaks, and transcription factor (TF) binding clusters are located and confined to the valleys immediately upstream of both peak 1 and 2 within the *Sox11* 3’UTR (Fig. 7B, B’2,B’5). In contrast, miRNA binding sites (which bind to 3’UTR sequences) show the opposite- and are completely confined to the peak1 and peak2 regions, never within valleys (Fig. 7B’3). Further, there is extremely high sequence conservation across species within *Sox11* 3’UTR peak 1 and 2, in fact conservation across these peaks is greater than across the *Sox11* CDS (Fig. 7B’4). This high degree of evolutionary conservation in 3’UTR sequence is true for some, but not all genes (Supp. Fig. 4).

We then assumed that i3’UTRs, as well as their particular peaks and valleys should be reflected in EST data-74 million stretches of RNA sequence generated from cDNA libraries; a sampling of the natural mRNA species. Of the 122 ESTs assigned to *Sox11*, 89% correspond to 3’UTR alone and contain no CDS sequence, with the rest (11%) corresponding to the CDS alone (Fig. 7B,B’1 red arrows, 3’UTR, green arrow, CDS). Strikingly, not one *Sox11* EST spans the 3’UTR-CDS junction even though 4% correspond to the 250 bp immediately upstream. Downstream after an EST gap for 750 bp there is an abrupt uptick with 9% of the ESTs corresponding to next 250 bp. There is not one EST that spans the 3’UTR valley between peak1 and 2 (Fig. 7 B’1; blue dotted circle). *Sox12* shows similar patterns, 84% of the Sox12 ESTs correspond to 3’UTR only, 5% to CDS only, and 10% span the junction (Fig. 7B). However, for other genes, for example *Nurr1*, most ESTs are uniquely CDS (Fig. 7B; green sidebar vs red sidebar). For others, in particular those with a Type (iii) pattern-a long 3’UTR with leading truncated CDS peak-there is good EST representation across the CDS-3’UTR junction, for example *Foxa2*, shows a high % of ESTs that cross the 3’UTR/CDS junction (Fig. 7B; right; yellow bar).

In summary for some genes we find very high 3’UTR conservation across species with gene sequence elements that are perfectly aligned perfectly with observed RNAseq peaks and valleys. For other genes, even though we observe highly differential 3’UTR/CDS expression by ISH (e.g. *Foxa2*), we do not find strict gene element and peak alignment. It may be for some genes, differential 3’UTR/CDS expression is not regulated from within the genome by gene elements, but is post-transcriptional.

## Discussion

We show here that in cells in all tissues, at all ages, and for many/all genes there is differential expression of coding regions (CDS) and parts or all of their 3’UTR sequences. This occurs in a non-random fashion, and within a tissue genes that show high 3’UTR/CDS expression are likely to be involved in the particular functions of that tissue, while genes with more equal CDS/3’UTR or > CDS/3’UTR involved in more general cell maintenance functions. In addition, 3’UTR/CDS ratios are not fixed but are dynamic and change as a cell changes state or developmental age. Within a 3’UTR we (and others) (1, 2), find non-continuous expression with peaks and valleys of RNA expressed-and with strikingly similar peaks and valley expression for many genes across tissues. For some genes, these peaks and valleys also show a correlation with miRNA and TF binding, as well as H3K27ac marks, suggesting that there may be elements intrinsic to the gene sequences that mediate the formation and the expression of isolated 3’UTRs. H3k27ac is known to be involved in increased expression of lncRNAs (29, 30). In addition, at least for some genes, the evolutionary conservation of these peaks suggests the importance and biological relevance of these isolated, expressed, 3’UTR sequences.

### Differential 3’UTR/CDS usage by ISH and cellular localization

The execution of multiple controls for our ISH procedure show it to be robust and sensitive, and the i3’UTR expression patterns allow interesting conjectures. By analogy to the known functions of lncRNAs, (i3’UTRs are by definition also lncRNAs), isolated 3’UTRs can bind to specific miRNAs and RNA binding proteins (RBPs) (31, 32), and likely able to add positive or negative value to the cognate protein’s expression and/or ability to function. One interesting example is Nurr1, while studies of Nurr1 protein show restricted expression, we find widespread *Nurr1* 3’UTR expression in both developing and adult tissues, often in the absence of high or even detectable levels of CDS. Genes that are thought to have very restricted expression based on their protein or CDS location may have widespread and high levels of 3’UTR expression in other cells. Nurr1 is one of three orphan-steroid receptors sub-family members; all suggested to be tumor suppressors (33-35), however their role can vary depending on cellular context (33). It is interesting to speculate that an interplay between isolated *Nurr1* 3’UTR levels and Nurr1 protein add to this contextualization. Here we showed high levels of *Nurr1* isolated 3’UTR sequences in proliferating cells (in the Ki67+ cells of the deep hair follicle) and high levels of *Nurr1* CDS in just adjacent Ki67-cells. Nurr1 is known to affect cell cycle (36), and it may be that highly expressed 3’UTR sequences in proliferating cells function in cell cycle dynamics, either together with their cognate or other proteins, or in the absence of protein.

Importantly we found many instances of isolated 3’UTR expression in the nucleus. Further, while proteins are often localized to cells with equal 3’UTR/CDS or high CDS, we find multiple notable exceptions; certain transcription factors are found to preferentially colocalize with their cognate i3’UTR sequences in the nucleus in some cells and some instances. While we cannot verify that CDS is entirely absent from these regions, what is clear is that there is less CDS than in nearby cells, thus we see “preferential’ localization of protein with i3’UTR sequences. While conjecture at this point, it may be that 3’UTR sequences in the nucleus stabilize/and or sequester cognate protein to delay degradation or temporarily prevent function. Studies to examine these questions are in progress. The specific localization of high levels of 3’UTR sequences to either the nucleus, or to axons, and the dynamic nature of these localizations have huge implications not only for nervous system axonal guidance synaptogenesis and plasticity, but for all cellular functions.

### Elusive gene families

Pinning down the precise or even general role of large gene families such as the *Sox* genes has been elusive due to imprecise knowledge about their gene and protein expression, as well as partial biological redundance, resulting in small phenotypes after gene deletion. Knowledge and manipulation of the distinct 3’UTR/CDS expression patterns of members of these gene families will provide us an additional handle from which to examine their function in neural and other development processes. Dual ISH of *Sox* mRNA expression reveals intricate patterns for each *Sox* gene in the nervous system with precise and abrupt changes in the use of the 3’UTR and/or the CDS during neuronal development. Extremely intriguing *Sox* gene 3’UTR/CDS patterns were observed in other tissues as well, but their description is beyond the scope of this article.

Further, the Netrin ligand-receptor family contains multiple distinct receptors with four Unc5 family receptors; with the particular function of the different Unc5 receptors somewhat indefinite in neuronal and vascular development as well as tumor suppression (37, 38). The very robust expression of *Unc5d* 3’UTR in blood vessels suggests that perhaps this alternate family member with highly expressed 3’UTR sequences may in some way enhance or modulate the specific known interactions of Ntn-1 and Unc5B in this tissue (39, 40).

### Stem cell biology

In two different stem cell niches we observed highly differential 3’UTR/CDS expression of the core pluripotent gene (PpG) network in progenitor cells. Importantly, in adjacent sections, non-PpG gens; (ie *Shh* and *Ascl1)* show more restricted patterns, and with more coincident 3’UTR/CDS expression. This is notable for two reasons, first PpG gene expression has not been highly documented outside of pluripotent stem cells. In agreement with other studies we find that PpG protein is low in these systems (not shown) however, their mRNA levels are robust and protein is reliably detected in a reasonable cohort of cells expressing the cognate mRNA. We expect that the precise regulation of the 3’UTR/CDS ratio, and thereby regulation of PpG protein expression might be critical for stem/progenitor cell function, possibly affecting the onset, duration and magnitude of proliferation and quiescence. PpG protein may be used sparingly, but cells appear to maintain specific 3’UTR/CDS expression levels, possibly to provide precise and titrated levels of protein. Secondly, we find abrupt changes in 3’UTR/CDS ratios in closely apposed but morphologically distinct tissues, such as *Sox2* and *Myc* in the whisker DP and a number of genes in Ki67+ vs – cells in the hair follicle. Sox2 is critical in DP development (26, 41) and shows equal 3’UTR/CDS in DP cells, whereas *Myc*, known for its role in proliferation, shows high 3’UTR. Interestingly, DP cells serve as a reservoir of stem cells (IPSC cells derive from the DP, (42-44); the high *Myc* 3’UTR-low CDS in these cells may allow them to maintain a non-proliferative, quiescent stem cell state.

### Dynamic 3’UTR/CDS

In addition to multiple genes that show specific 3’UTR and CDS expression, these ratios are dynamic and depend on the cell state-we show they change across development (2), after stress, or with proliferative challenge.

While some genes showed prominent localization of their 3’UTR sequences to P bodies, marked by DDX6, or the nuclear membrane marked by Lamin, this was not consistent across genes and or cell types and does not appear to be requisite. However we never observed isolated CDS co-localized with these markers. Examination of individual i3’UTR sequences with these structures in particular contexts will be necessary and valuable.

### Bioinformatics

This bioinformatic study is the first to systematically examine gene classes that are likely to express high 3UTR/CDS ratios or the reverse, high CDS compared to 3’UTR. They confirm and extend previous findings from our work and others using RNAseq and ISH (1-3). Here, from random select open source RNAseq data sets we find that differential 3’UTR/CDS expression is commonly observed, appears highly regulated-both in the expression of either the 3’UTR or CDS and in the use of particular 3’UTR regions. We further show that although isolated 3’UTR sequences are not always contiguous with the CDS, nor are they continuous along the length of the 3’UTR, their “peaks” and “valleys”-at least for some genes such as *Sox11* and *12* align perfectly with various gene elements. For both genes the 3’UTRs are highly conserved across species, with the peaks preceded by TF clusters, DNase1 hypersensitive sites, Cpg islands and H3K27ac marks-all hallmarks of gene transcription initiation. Further and in contrast, known miRNA binding sites align perfectly with the RNAseq peaks and not with the valleys. Finally, the library of known ESTs aligns exactly with the peaks and not with the valleys. Both *Sox11* and *12* have few exons and long 3’UTRs. For other genes, even though we see highly differential 3’UTR/CDS expression by in situ hybridization and RNAseq, these elements may be, but are not necessarily as perfectly aligned. It is possible that for some genes, the expression of isolated 3’UTR sequences is genome driven, while for others post-translational mechanisms are important. Further bioinformatic studies will shed light on these possibilities. Again, because it appears to be gene dependent, we provide examples of “types” of genes observed.

Finally, by examining the GO categories of high CDS and high 3’UTR genes across multiple tissues we find that high 3’UTR genes are tissue specific-related to the particular functions of, or the developmental state of each tissue. In contrast the bulk of significant GO categories for high CDS genes are related to cell maintenance. However, in testis there were also significantly GO categories for high CDS genes that were tissue specific. Our interpretation of this data is that cells use high isolated 3’UTR expression for tissue specific genes when the cell is in a metastable state, either developmentally or when carrying out its specific function (ie gtpase signaling in adult heart). We are very interested in the possibility that conversely, the identification of i3’UTR genes can inform us about a tissue at hand, either in chronic (cancer) or acute (viral infection) disease states.

In sum, together this data shows there is a strong proclivity for high 3’UTR/CDS in cells and for genes involved in transition states, during both proliferative and developmental maturation events.

## Materials and Methods

### Fluorescence in situ hybridization (ISH) and immunochemistry

Two color fluorescence ISH was performed as previously described (2), using the TSA Plus Cyanine 3 & Fluorescein (NEL753001KT, PerkinElmer, Waltham, MA) according to the manufacturer’s instructions. Briefly, embryonic and adult tissues, or cells in culture were fixed with 4% paraformaldehyde (PFA) in 0.1 M phosphate buffer (PB). Tissues were cryoprotected, embedded in OCT and stored at −80C until 16 μm cryostat sections were collected (Lecia, Buffalo Grove, IL) onto Superfrost plus glass slides (Invitrogen). For both tissues and cells, after post-fixation in 4% PFA/PB, Proteinase K treatment, acetylation (1% triethanolamine and 0.25% acetic anhydride and prehybridization (50% formamide, 5X SSC, 5X Denhardt’s solution, 0.5 mg/ml Herring sperm DNA and 250 μg/ml Yeast tRNA) at room temperature for 1 hour, tissue sections were hybridized overnight (cells for 3 hours) with fluorescein labeled CDS and Digoxygenin (DIG) labeled probes 3’UTR probes at 56°C. Specimens were then sequentially post-stained with fluorescein or Cy3 chromogens, respectively, and the corresponding Alexa Fluor 647 secondary antibodies and Hoechst 33342 for DNA staining, and mounted with Mounting Medium. Antibodies in Supp. Table 1. In situ probes in Supp. Table 2.

### Cell culture

NIH3T3 cells were grown at 37^’^C in DMEM, 10% FBS and 0.5 mg/ml Penicillin Streptomycin Glutamine. Sodium azide (Sigma/Aldrich) was added to cell media 30 min before fixation.

### Micropattern cell culture

Micropatten CYTOO (Arena A, CYTOO, France) chip cell culture was performed as previously described (Morgani S. et. al.). Patterns of 800, 500, 225, 120 and 80 um circles are dispersed over the full surface, with 25, 144, 576, 900 and 1296 colonies of each size respectively. Briefly, 700 ul drops of 20 mg/ml Laminin (L20202)(Sigma) in PBS without calcium and magnesium (PBS-/-) in were spotted onto parafilm lain in a 15 cm tissue culture plate. After one wash in PBS-/-, chips were inverted on top of the drops for 2h at 37 °C. They were then washed 5x with PBS-/- and a single cell suspension of 3T3 cells (2 × 10^6^) was evenly plated onto chips lain in 6-well plates for two days in complete DMEM medium before fixation and ISH.

### Bioinformatic analyses

Raw sequencing reads were acquired from GEO and ENCODE as follows; Fig. 5; differentiated tissue for GO enrichment analysis GSM2493731, Fig. 6; ES cell, ENCSR000CWC, E11 forebrain, ENCSR160IIN, E15 midbrain, ENCSR557RMA, E15 heart, ENCSR597UZW, P0 hindbrain, ENCSR017JEG, adult cortex, ENCSR000BZS, P0 midbrain, ENCSR719NAJ, P0 skeletal muscle, ENCSR946HWC, E15 DA neurons, GSE75596. All were aligned to the mouse genome (mm10) using STAR (v2.7.3a) as default parameters. Mapped files were filtered with minimum mapping quality 20 using samtools (v1.11) and CDS/UTR expression quantification analysis was performed for all samples using bedtools/coveragebed (v2.17.0) and normalized to library size. For each gene, we quantified the relative abundance of 3’UTR to CDS is quantified and expressed as the fractional ratio of (3’UTR mean coverage)/(3’UTR mean coverage + CDS mean coverage) which ranges from 0 highest CDS/3’UTR to 1, highest 3’UTR/CDS. Gene ontology (GO) enrichment analysis was performed using the Metascape online tool (http://metascape.org/gp/index.html) (Tripathi et al., 2015), and the top GO categories were selected according to the binomial P values.

## Acknowledgments

We thank Saranya Santoosh Kumar and Melanie O Rourke cell culture work, Dr. Hudspeth for the Sox11 antibody, Dr. Lei (University of New England) for the Sox11 KO mice, Susan Morton for the rabbit Foxa2 antibodies and Dr. S. Linnnarssen for bioinformatic discussion.

## Notes

### Competing Interest Statement

The authors have declared no competing interest.

### Summary of Updates

Figure resolution, author order

