## Supplementary material for "Distinct expression of select and transcriptome-wide isolated 3’UTRs suggests critical roles in development and transition states": Supp. figs

### Ji et al Supporting information

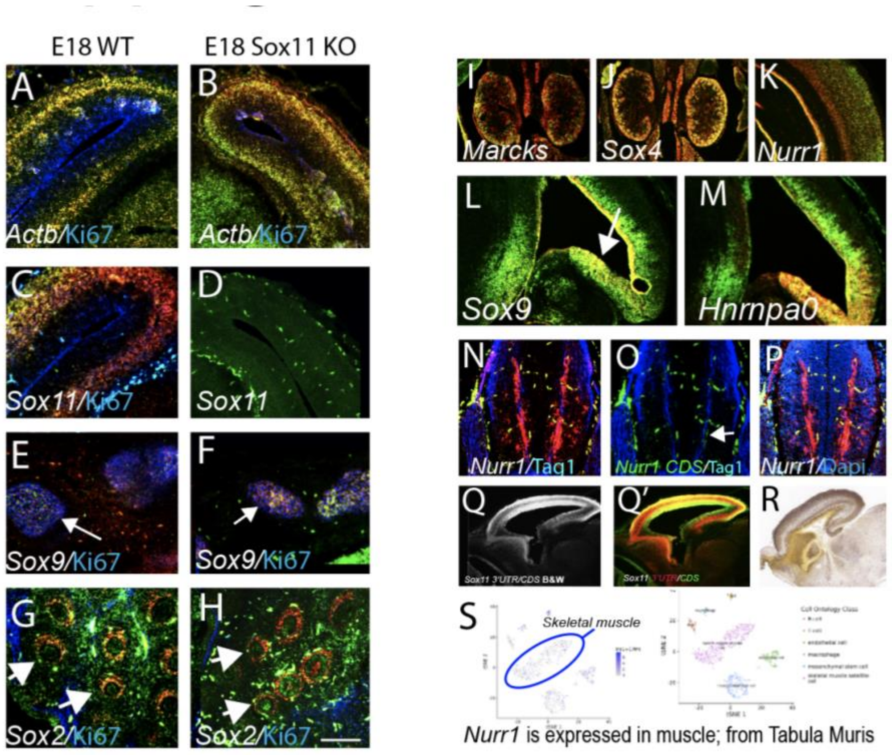


**S1 Fig. As relates to Fig 1.** (A-H) In situ hybridization for various probes in WT E18 and *Sox11* KO E18 tissues. All probes show signal in the *Sox11* KO animals except *Sox11* (green spots are background; compare C and D). (I, J) lower power views of Fig 1D,E showing entire kidney. (K) *Nurr1* in adult brain. (L-M) adjacent sections to Fig. 1 K-N, showing additional genes. (N-P) E11 spinal cord sections showing *Nurr1* 3'UTR/CDS (N) with low, but not "no signal" for the *Nurr1* CDS (O; green spots are background, white arrow points to signal). (P) same section with DAPI stain to show that background is not cellular and that many cells are *Nurr1* CDS and 3'UTR negative. (Q-R'), (Q,Q') *Sox11* 3'UTR/CDS in E15 sagittal brain in black and white (Q) and color (Q') compared to similar photo using the Allen brain Atlas CDS probe (R). With only one probe (Q,R) a meaningful pattern of *Sox11* is difficult to discern while the dual 3'UTR/CDS pattern suggests developmentally relevant expression (Q'). (S) Tabula muris data (https://tabula-muris.ds.czbiohub.org) showing widespread expression of *Nurr1* outside the brain and dopaminergic neurons. Highlighted is skeletal muscle. Scale bar, A-D; 500, E,F; 200, G,H; 400, I,J; 1500, K; 700, L,M; 300, N-P; 160,Q-R; 522 um.

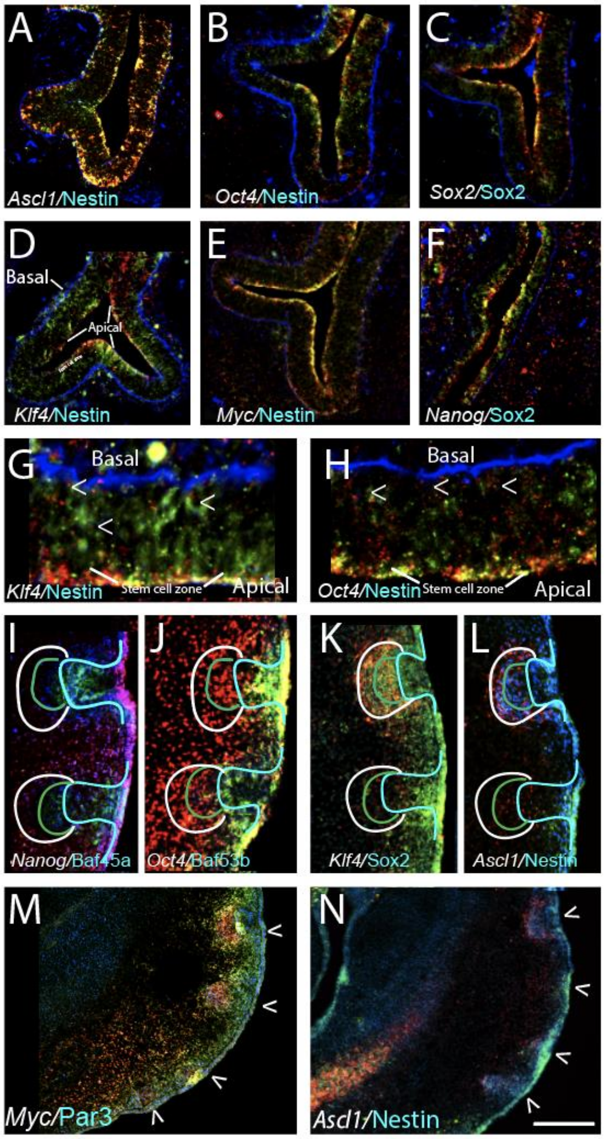


**S2 Fig. As relates to 2.** (A-F) Adjacent E12 OlfE sections to those in Fig. 2B,C, showing additional genes. All PpGs are expressed. (G,H) sections corresponding to Fig. 2D-G with additional PpG genes, Klf4 andOct4. (M,N) adjacent sections to Fig. 2 I,J with additional, indicated genes. Scale bar, A-C; 200, G-H; 150, I-L; 55, M,N; 350 um.

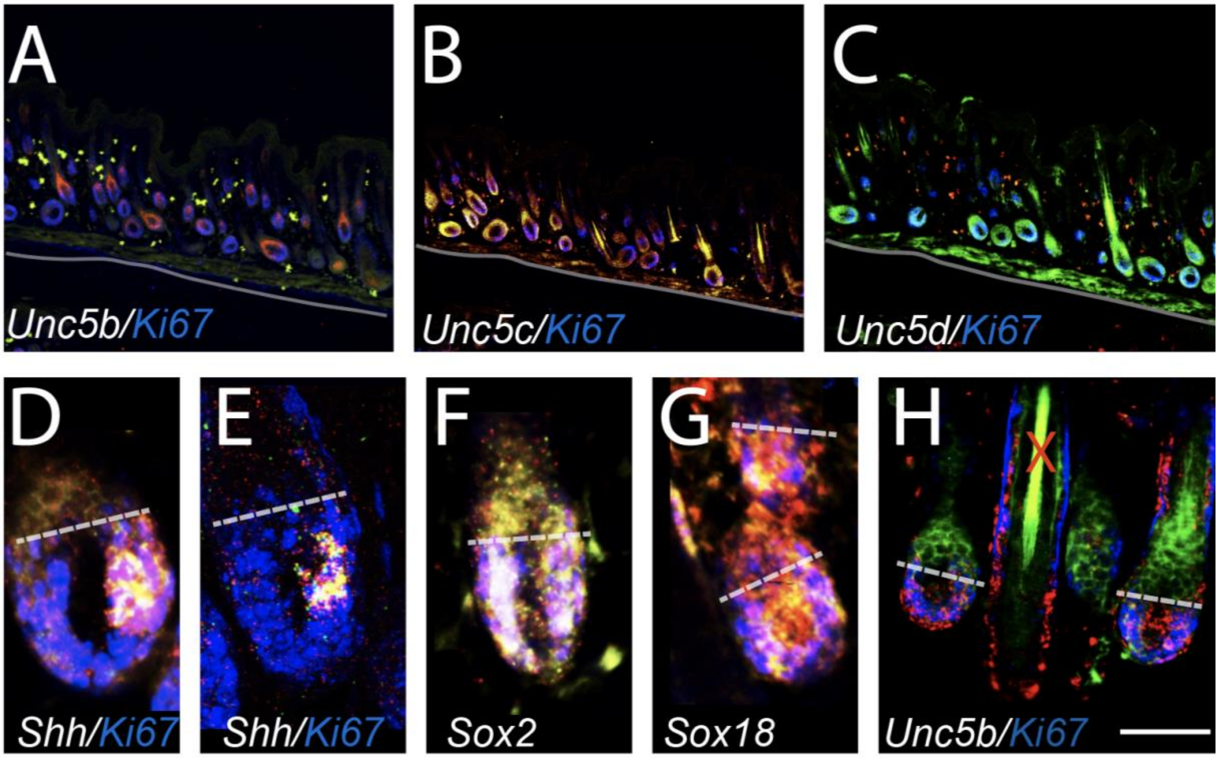


**S3 Fig. As relates to Fig. 3.** (A-C) Skin sections adjacent to Fig. 3D-F, showing additional genes as indicated. (D-G), adjacent sections to Fig. 3G-Q, showing additional genes as indicated. (D) *Shh* expression in Ki67+ cells (E) same as D but confocal image, note that pattern is identical in D,E. (H) same image as Fig. 2 M, M' but with hair shaft not cropped; note high auto-fluorescence in hair shaft, marked by red X. Scale bar A-C; 700, D-G; 85, H; 110 um.

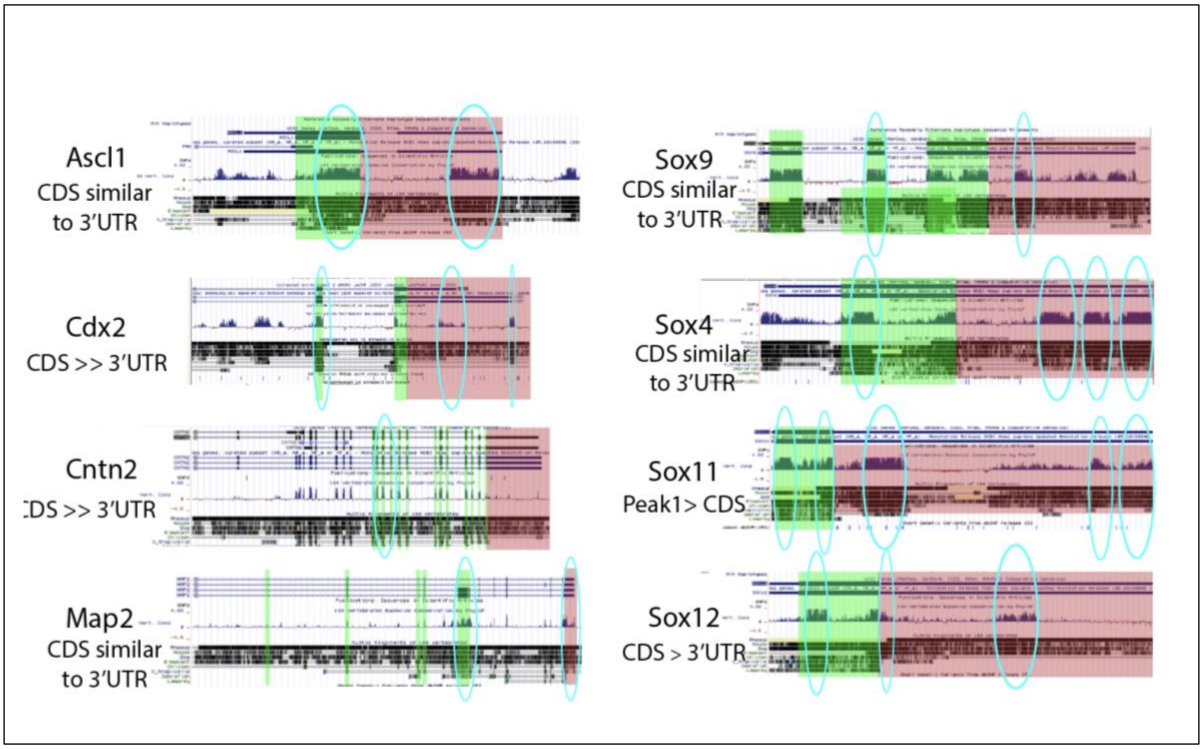


**S4 Fig. As relates to Fig. 7. Conservation and ESTs.** Some genes do not show high conservation within their 3'UTR across species, ie Cntn2 and Cdx2. Red shaded areas are 3'UTR, green CDS. Blue circles show regions of conservation.

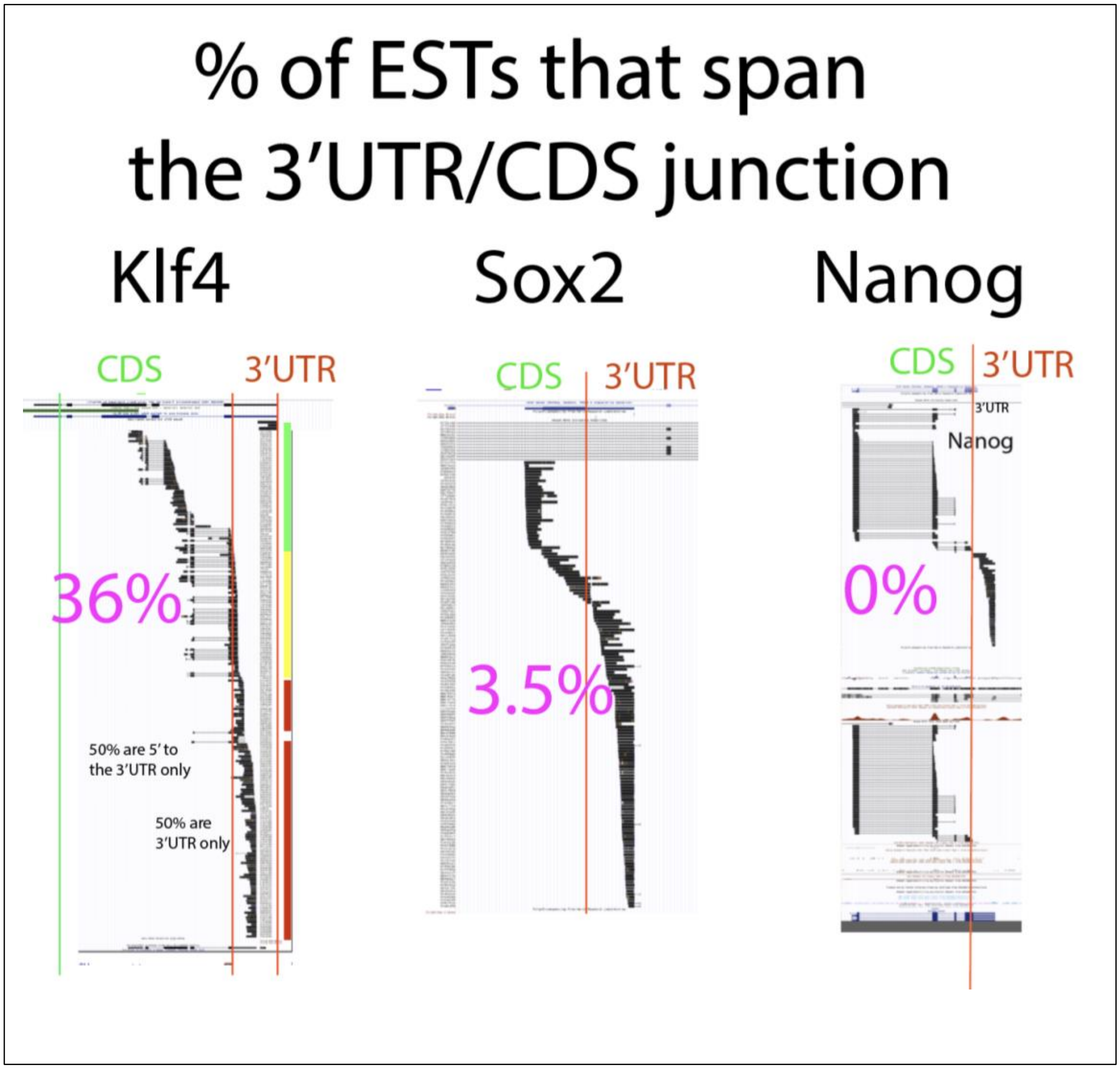


**S5 Fig. As relates to Fig7. Percent of EST's that span the 3'UTR/CDS junction for three additional genes.** *Klf4* shows a high preponderance of ESTs that span the 3'UTR/CDS junction, while the percent for *Sox2* is very low, and not one EST spans the junction for *Nanog*.

| Ab | antigen | Dilution | Cat. # | Company |
|---|---|---|---|---|
| Tag1 | Axon guidance molecule | 1:500 | AF4439 | R&D Systems |
| Nestin | Neural stem | 1:1000 | Ab22035 | Abcam |
| Sox2 | Pluripotent gene | 1:1000 | | R&D Systems |
| Baf45a | SWI/SNF complex Neural stem cells | 1:30 | N/A | Gift Dr. G. Crabtree |
| Baf53b | SWI/SNF complex; more differentiated cells | 1:30 | | Gift Dr. G. Crabtree |
| MAP2 | Structural protein | 1:1000 | NB300-213 | Novus Biologicals |
| Foxa2 mouse<br><br>4C7 against chick | Transcription factor in DA neurons | 1:50 | N/A | Gift Susan Morton |
| Foxa2 K2 rabbit anti hnf3b | Transcription factor in DA neurons | 1:10000 | N/A | Gift Susan Morton |
| Ki67 | Marker of active proliferation | 1:1000 | | Abcam |
| Ddx6 | P-body marker | 1:100 | Pa5-55012 | Invitrogen |
| Xrn1 | RNA degradation path; localized to P-bodies | 1:100 | Pa5 57110 | Invitrogen |
| Rcd8 | Mrna decapping P-body marker | 1:50 | Pa530485 | Invitrogen |

| | | | | |
|---|---|---|---|---|
| Lamin | Nuclear membrane | 1:100 | 33-2100 | Invitrogen |

**Supp. Table 1.** Antibody list; companies, catalog number and dilution for antibodies used in these studies.

**Supp. Table 2.** In situ probe sequences
