## Supplementary material for "Distinct expression of select and transcriptome-wide isolated 3’UTRs suggests critical roles in development and transition states": Supp table 2

| Supp Table2 | | | |
|---|---|---|---|
| Gene | CDS | 3’UTR | NCBI mRNA link |
| Sox9 | AGGAAGCTGGCAGACCAGTACCC<br>GCATCTGCACAACGCGGAGCTCAG<br>CAAGACTCTGGGCAAGCTCTGGAG<br>GCTGCTGAACGAGAGCGAGAAGA<br>GACCCTTCGTGGAGGAGGCGGAG<br>CGGCTGCGCGTGCAGCACAAGAA<br>AGACCACCCCGATTACAAGTACCA<br>GCCCCGGCGGAGGAAGTCGGTGA<br>AGAACGGACAAGCGGAGGCCGAA<br>GAGGCCACGGAACAGACTCACATC<br>TCTCCTAATGCTATCTTCAAGGCGC<br>TGCAAGCCGACTCCCCACATTCCTC<br>CTCCGGCATGAGTGAGGTGCACTC<br>CCCGGGCGAGCACTCTGGGCAATC<br>TCAGGGTCCGCCGACCCCACCCAC<br>CACTCCCAAAACCGACGTGCAAGC<br>TGGCAAAGTTGATCTGAAGCGAGA<br>GGG | AAGTGTGTGTGCCGTGGATAGCCC<br>CTTGGCTGCTCTCCTGCAGAGAGA<br>CATCGGACAGACCTTAATTCTTACT<br>CACTGCTGTGGCTGGAGAGTATAA<br>GGAATGCTTTTTCTTTTTTTCTTTCT<br>TTCTTTCTTTTTTTTTTTTAAGACAG<br>CAGTCTTTTTTTTTAATTTAAAAAA<br>AAAAAGATATATTAACAGTTTTAG<br>AAGTCAGTAGAATAAAACCTTAAA<br>GCGTTCTTATAATATGGCATCTTTC<br>GATTTCTGTATAAAAACAGACCTTT<br>TAAAAAATATTTCTGTAACTTAAGA<br>AACCTGACATTTATGTCATATTTTC<br>TCTTTAGGTAAGATTTGGTTTGTTT<br>GTGTCTTGTTTGTTTCCCTCTCCAA<br>ATTCTTCCTTCTTTGTGCACCCTGCC<br>TTTCTTCCCTTCCATCCT | https://www.ncbi.nlm.nih.gov/nuccore/NM_011448.4 |
| Sox11 | CTTCATGGTGTGGTCC<br>AAGATCGAGCGCAGGAAGATCAT<br>GGAGCAGTCGCCCGACATGCACAA<br>CGCCGAGATCTCCAAGAGGCTGG<br>GCAAGCGCTGGAAGATGCTGAAG<br>GACAGCGAGAAGATCCCGTTCATC<br>AGGGAGGCGGAGCGCCTGCGCCT<br>CAAGCACATGGCTGATTATCCCGA<br>CTACAAGTACCGGCCGCGCAAAAA<br>GCCCAAGACGGACCCAGCGGCC<br>AAGCCCAGCGCGGGCCAGAGCCC<br>CGACAAGAGCGCGGCGGGCGCCA<br>AGGCAGCCAAGGGCCCCGGCAAG<br>AAGTGCGCCAAGCTCAAGGCGCCT<br>GCGGGCAAGGCGGGCGCGGGCAA<br>GGCGGCGCAGCCGGGGGACTGCG<br>CCGCGGGCAAGGCAGCCAAGTGC<br>GTCTTCCTGGACGACGACGATGAA<br>GACGACGACGAAGATG | CTGGCCAGGCTTCAATAAAAGGTT<br>AGCTGCATTTTTTTTCCTCCTAAGT<br>AAGTCTTATTTTTAACTGAGCATTG<br>ACAGTATCTTAAAATGGTAACGTG<br>GGCGTGGGCGTTGTGTGCATAGCA<br>GTCTAGCCGGTTGGGTACCCTGCT<br>CCTGTACCTAGTTCACAGACTCGA<br>GTGCATTTTTTTTTGGCGAGATTTC<br>CATCTTTGAAGAAACAAAAAAACA<br>AAAACACCTGAACCAGCGTTCTCTT<br>ATTCTTTAAGCTGTGGAAATAATTT<br>CCAGTTTCTACATTCTCGATATGCA<br>TCCTTATTAAAAAGATAATACGAAT<br>GAAAGGCAGTGTGCTTAAAGTGTG<br>CTTTGCAAATACATGTTATGAATGA<br>CTACGGTCACTGGGCAAATT | https://www.ncbi.nlm.nih.gov/nuccore/NM_009234.6 |
| Marcks | AGGAGAATGGCCACGTAAAAGTG<br>AACGGGGACGCGTCTCCCGCCGCC<br>GCCGAGCCGGGCGCCAAGGAGG<br>AGCTGCAAGCCAACGGCAGCGCCC<br>CGGCCGCCGACAAGGAGGAGCCC<br>GCGAGCGGCAGTGCCGCGACCCC<br>CGCCGCGGCCGAAAAGGATGAGG<br>CTGCCGCGGCCACCGAGCCGGGC<br>GCCGGCGCGGCCGACAAGGAGGC<br>TGCGGAGGCCGAGCCCGCCGAGC<br>CCAGCTCCCCGGCCGCCGAGGCCG<br>AGGGCGCGTCCGCCTCCTCCACGT<br>CGTCGCCCAAGGCGGAGGACGGG<br>GCCGCGCCGTCGCCCAGCAGCGAG<br>ACCCCGAAAAAAAAAAAAGAAGCG<br>CTTTTCCTTCAAGAAGTCCTTCAAG | TTGTGAGGCAGGTTTACAACACTA<br>CACGTTTTGAATAAGAAGGAA<br>AGAGAAAAAAATAAAAACACCAAT<br>ACCCAGATTTAAAAAAAAAAAAAA<br>AGATCATAGTCTTAGGAGTTCA<br>TGTAAACCATAGGAACTTCCTGCTT<br>ATCTCATGTTAGCTGTACCAGTCAG<br>TGATTAAGTAGAACTACAAG<br>TTGTATAGGCTTTATTGTTTATTGC<br>TGGTTTATGACCTTAATAAAGTGTA<br>ATTATGTATTACCAGCAGGG<br>TGTTTTTAACTGTGACTATTGTATA<br>AAAACAAATCTTGATATCCTTCAGA<br>AGCACATGAAGTTTGCAAGT<br>CTCCACCCTGCCCATTTTTGTAAAA<br>CTGCAGTCATCTTGGACCTTTTAAA | https://www.ncbi.nlm.nih.gov/nuccore/NM_008538.2 |

| | | | |
|---|---|---|---|
| | CTGAGCGGCTTCTCCTTCAAGAAG AGCAAGAAGGAGTCGG | CACAAAATTTTAAACTCAAC CAAGCTGTGATACGTGGAA | |
| Sox4 | GCCCAAGAAGAGCTGTGGCCCCAA GGTGGCGGGCAGCTCGGTCGGCA AGCCCCACGCTAAGCTGGTCCCGG CGGGCGGCAGCAAGGCGGCTGCA TCGTTCTCTCCAGAGCAAGCTGCCC TGCTGCCCCTGGGGGAGCCCACGG CCGTCTACAAGGTGCGGACTCCCA GTGCGGCCACTCCGGCCGCCTCCT CCTCGCCGTCCAGTGCGCTGGCCA CCCCAGCCAAACACCCTGCCGACA AGAAAGTGAAGCGCGTCTACCTGT TTGGAAGCCTGGGCGCTTCGGCGT CTCCCGTCGGGGGCCTGGGAGCG AGCGCCGACCCCAGTGATCCACTG GGGTTGTACGAAGATGGAGGC | GCCTCATGGTCAAGAAAGGAGGG GGAAAATCCAGCGTGCCCCATCTC CTACCCACCCCCTCTTTGTATTCTCT TTGTATTTTTCCCCTTCTTAAAATTT CTTTTTCTGCAATGAAGACAGAAA GAAGGCTCTGGGGTGATGCGTTTG GCATTTGTGTTGAGCTTAGGGGAG CATTGGCATGGAGAAACTCCACGC TGGCGAAGTCCCGGGCTGGGCTTT CTCCTCCTCCCCCCACCTTTTTTCCC CTTGTCCTTGCAGCTGGAGATGTG CTGGGAGTAGCAGGCCAGCCTCG GAAATGGACATGGGGACCTCGTG GAAGCCACAGCAACCTGGTTGGG GATGCAGAAGGAC | https://www.ncbi.nlm.nih.gov/nuccore/NM_009238.3 |
| Nurr1 | GGACGATCCGGGCTCCCTTCACAA CTTCCACCAGAACTACGTGGCCACT ACGCATATGATCGAGCAGAGGAA GACACCTGTCTCCCGCCTGTCACTC TTCTCCTTTAAGCAGTCGCCCCCGG GCACTCCTGTGTCTAGCTGCCAGAT GCGCTTCGACGGGCCTCTGCACGT CCCCATGAACCCGGAGCCCGCGGG CAGCCACCACGTAGTGGATGGGCA GACCTTCGCCGTGCCCAACCCCATT CGCAAGCCGGCATCCATGGGCTTC CCGGGCCTGCAGATCGGCCACGCA TCGCAGTTGCTTGACACGCAGGTG CCCTCGCCGCCGTCCCGGGGCTCT CCCTCCAATGAGGGTCTGTGCGCT GTTTGCGGTGACAACGCGGCCTGT CAGCACTACGGTGTTCGCACTTGT GAGGGCT | AAACAAACCACAAATAAAAACTGT CGCTATTTCCTAACCTGCAGGCAG AACCTGAAAGGGCATTTTGGCTCC GGGGCATCCTGGATTTAGAAAACG GACAGCACACAGTACAGTGGTATA AACTTTTTATTATCAGTTCAAAATC AGTTTGTTGTTCAGAAGAAAGATT GCTAATGTATGATGGGAAATGTTT GGCCATGCTTGCTTGTTGCAGTTAA GACAAATGTAACACAC | https://www.ncbi.nlm.nih.gov/nuccore/NM_013613.2 |
| Actb | GGGAATGGGTCAGAAGGACTCCT ATGTGGGTGACGAG | GTGGCTGAGGACTTTGTACATTGT TTTGTTTTTTTTTTTTTTTTGGTTTT | https://www.ncbi.nlm.nih.gov/nuccore/NM_007393.5 |

| | | | |
|---|---|---|---|
| | GCCCAGAGCAAGAGAGGTATCCTG<br>ACCCTGAAGTACCCCATTGAACAT<br>GGCATTGTTACCAACTGGGACG<br>ACATGGAGAAGATCTGGCACCACA<br>CCTTCTACAATGAGCTGCGTGTGG<br>CCCCTGAGGAGCACCCTGTGCT<br>GCTCACCGAGGCCCCCCTGAACCC<br>TAAGGCCAACCGTGAAAAGATGAC<br>CCAGATCATGTTTGAGACCTTC<br>AACACCCCAGCCATGTACGTAGCC<br>ATCCAGGCTGTGCTGTCCCTGTATG<br>CCTCTGGTCGTACCACAGGCA<br>TTGTGATGGACTCCGGAGA | GTCTTTTTTTAATAGTCATTCCAAG<br>TATCCATGAAATAAGTGGTTACAG<br>GAAGTCCCTCACCCTCCCAAA<br>AGCCACCCCCACTCCTAAGAGGAG<br>GATGGTCGCGTCCATGCCCTGAGT<br>CCACCCCGGGGAAGGTGACAGC<br>ATTGCTTCTGTGTAAATTATGTACT<br>GCAAAAATTTTTTTAAATCTTCCGC<br>CTTAATACTTCATTTTTGTT<br>TTTAATTTCTGAATGGCCCAGGTCT<br>GAGGCCTCCCTTTTTTTTGTCCCCC<br>CAACTTGATGTATGAAGGCT<br>TTGGTCTCCC | |
| Sox12 | CACGAGCGGAGAAAAATCATGGA<br>CCAGTGGCCCGACATGCACAACGC<br>TGAGATCTCCAAGCGCCTGGGCCG<br>CCGCTGGCAGCTGCTGCAGGACTC<br>GGAGAAGATCCCGTTCGTGCGGG<br>AGGCGGAGCGGCTGCGCCTCAAG<br>CACATGGCGGACTACCCGGACTAC<br>AAGTACCGGCCTCGCAAGAAGAGC<br>AAGGGGG<br>CGCCCGCCAAGGCGCGGCCCCGCC<br>CCCCCGGAGGCGGCGGTGGTGGC<br>AGTCGGCTGAAACCCGGGCCACA<br>GCTGCCGGGCCGCGGGGGCCGCC<br>GAGCGTCGGGAGGACCTCTGGGG<br>GGCGGCGCGGCGGCGCCGGAGGA<br>CGACGACGAAGACGAAGAGGAG | GGC<br>TCCTCCCTAAGTCCATCACCCCTTC<br>CTATCACTCAAATCCAGTCTCAGCA<br>ACCACAACTGCAGTATTCAG<br>TGGGAGAAACTGAAGCACGTTGTT<br>GCTAGGGTTTAGGACAGGCATACC<br>AGCAGCCGGGTGGCAAGAATGA<br>GGACCATGTCTAAACAAACTTGGC<br>TTGAGGAGAGACCAGGCCAGGGG<br>ACGGTGGCAGAGCTAGTAGTGGA<br>CTTGGGTGCTTTGCACCTCGGTCTG<br>CCCTTGTGAGAAATGGGGTGGCAC<br>TGCTGTAGTTAGGAAGCTGTG<br>TTGAGATTAGGTTGCCAAGTCTGC<br>TCCACTGGTCTCCCGCTCCAGCCCC<br>GCCCCTCCAGCCTCCCCATCC<br>TCCCTCTGCCCTCACTACCTGTATCT<br>CACCGGCG | https://www.ncbi.nlm.nih.gov/nuccore/NM_011438.2 |
| Map2 | TGGTGGACGTGTGAAAATTGAGA<br>GTGTAAAACTGGATTTCAAGGAAA<br>AGGCCCAAGCTAAAGTT<br>GGCTCACTTGACAATGCTCACCAC<br>GTACCTGGAGGTGGTAATGTGAAG<br>ATTGACAGCCAAAAGTTGAACT<br>TCAGAGAGCATGCAAAGGCCCGG<br>GTAGATCACGGGGCTGAGATCATC<br>ACACAGTCCCCAAGCAGGTCCAG<br>CGTGGCATCACCCCGACGACTCAG<br>CAACGTCTCATCTTCTGGAAGCATC<br>AACCTGCTCGAATCCCCTCAG<br>CTTGCCACTTTGGCTGAGGATGTC<br>ACTG | ATTGCAGGAAGGAAAGAGCATGT<br>AAGAAACACATTTTTTAAAGTGTTA<br>TTTTGTATAAATGGGAAGAAA<br>GACGCAATTAAGTTATTGACACTT<br>GGGACCTGGACGAGTATATCAGA<br>GTATGCCATTCCAATAAATTATT<br>GAACTACAAGCTAGATTTAAGGCA<br>TTTGAGCGTTGGTTGAAGAAGTGG<br>TGTCAAAGTGCATCTCTTAGGA<br>TTGATGCACTTTTGTTAGGATGGG<br>CTTGTGTCTGATTAGAATGTCAGTC<br>GATTGGCTAGATTTATATCCA<br>CACAATCAGTTTCACACCCCCATTC<br>CATCTGTTTGATACAGTATTATAGA<br>TATAAATATATATATATATT<br>TCTCTGTGGCCATTTGTGA | https://www.ncbi.nlm.nih.gov/nuccore/NM_001039934.1 |
| Ascl1 | GTGGCCGACAGCCAGCCCTCAGGG<br>GGCGGTCACAAGTCAGCGGCCAA<br>GCAGGTCAAGCGCCAGCGCTCGTC<br>CTCTCCGGAACTGATGCGCTGCAA<br>ACGCCGGCTCAACTTCAGCGGCTT<br>CGGCTACAGCCTGCCACAGCAGCA<br>GCCGGCCGCCGTGGCGCGCCGCA | AGACAGACACTATATTAACTCCCAA<br>CCACTAACAGGCAGGGCTGGAAG<br>CGCGCATGTGCAAGTGCCTTCACC<br>TCCCACTCTCTGTCAGAGCTGTCTT<br>AGCCCCCTGAAACTGGGTTGATGT<br>CTTTCCTCAGTCACCCCCATTCCAG<br>CGATCTATGGACATTTGCCTCCATT | https://www.ncbi.nlm.nih.gov/nuccore/NM_008553.5 |

| | | | |
|---|---|---|---|
| | ACGAGCGCGAGCGCAACCGGGTC<br>AAGTTGGTCAACCTGGGTTTTGCC<br>ACCCTCCGGGAGCATGTCCCCAAC<br>GGCGCGGCCAACAAGAAGATGAG<br>CAAGGTGGAGACGCTGCGCTCGG<br>CGGTCGAGTACATCCGCGCGCTGC<br>AGCAGCTGCTGGACGAGCACGAC<br>GCGGTGAGCGCTGCCTTTCAGGCG<br>GGCGTCCTGTCGCCCACCATCTCCC<br>CCAACTACTCCAACGACTTG | GAAGCAACGTCAGTTCTCGGACAG<br>CCTTTCCCTCTCCTGGTGGCCTCCT<br>CCCCAAACCCCACATCGCCCTCCCA<br>CGGTCTTTGCTTCTGTTTTCTTCATA<br>GAATGCTTCCAATCTTTGTGAATTT<br>TTTTATTATAAGAAAAAAATCTATT<br>TGTATCTATCCTAACCAGTTTGGGG<br>ATATATTAAGATATTTTTGTACATA<br>AGAAAAAGAGAGAGA | |
| Myc | GTCGCTACGTCCTTCTCCCCAAGGG<br>AAGACGATGACGGCGGCGGTGGC<br>AACTTCTCCACCGCCGATCAGCTG<br>GAGATGATGACCGAGTTACTTGGA<br>GGAGACATGGTGAACCAGAGCTTC<br>ATCTGCGATCCTGACGACGAGACC<br>TTCATCAAGAACATCATCATCCAGG<br>ACTGTATGTGGAGCGGTTTCTCAG<br>CCGCTGCCAAGCTGGTCTCGGAGA<br>AGCTGGCCTCCTACCAGGCTGCGC<br>GCAAAGACAGCACCAGCCTGAGCC<br>CCGCCCGCGGGCACAGCGTCTGCT<br>CCACCTCCAGCCTGTACCTGCAGG<br>ACCTCACCGCCGCCGCGTCCGAGT<br>GCATTGACCCCTCAGTGGTCTTTCC<br>CTACCCGCTCAACGACAGCAGCTC<br>GCCCAAATCCTGTACCTCGTCCGAT<br>TCCACGGCCT | ACTGACCTAACTCGAGGAGGAGCT<br>GGAATCTCTCGTGAGAGTAAGGAG<br>AACGGTTCCTTCTGACAGAACTGA<br>TGCGCTGGAATTAAAATGCATGCT<br>CAAAGCCTAACCTCACAACCTTGG<br>CTGGGGCTTTGGGACTGTAAGCTT<br>CAGCCATAATTTTAACTGCCTCAAA<br>CTTAAATAGTATAAAAGAACTTTTT<br>TTTATGCTTCCCATCTTTTTTCTTTTT<br>CCTTTTAACAGATTTGTATTTAATT<br>GTTTTTTTAAAAAAATCTTAAAATC<br>TATCCAATTTTCCCATGTAAATAGG<br>GCCTTGAAATGTAAATAACTTTAAT<br>AAAACGTTTATAACAGTTACAAAA<br>GA | https://www.ncbi.nlm.nih.gov/nuccore/NM_010849.4 |
| Nanog | CCCACAGTTTGCCTAGTTCTGAGG<br>AAGCATCGAATTCTGGGAACGCCT<br>CATCAATGCCTGCAGTTTTTCATCC<br>CGAGAACTATTCTTGCTTACAAGG<br>GTCTGCTACTGAGATGCTCTGCAC<br>AGAGGCTGCCTCTCCTCGCCCTTCC<br>TCTGAAGACCTGCCTCTTCAAGGC<br>AGCCCTGATTCTTCTACCAGTCCCA<br>AACAAAAGCTCTCAAGTCCTGAGG<br>CTGACAAGGGCCCTGAGGAGGAG<br>GAGAACAAGGTCCTTGCCAGGAA<br>GCAGAAGATGCGGACTGTGTTCTC<br>TCAGGCCCAGCTGTGTGCACTCAA<br>GGACAGGTTTCAGAAGCAGAAGT<br>ACCTCAGCCTCCAGCAGATGCAAG<br>AACTCTCCTCCATTCTGAACCTGAG<br>CTATAAGCAGGTTAA | GGAGACAGTG<br>AGGTGCATATACTCTCTCCTTCCCA<br>AGAATAAGTGCTTGAACACCCTTA<br>CCCACGCCCACCCACCCATGC<br>TAGTCTTTTTTCTTAGAAGCGTGGG<br>TCTTGGTATACACTGTGTCATTTTG<br>AGGGGTGAGGTTTAAAAGTA<br>TATACAAAGTATAACGATATGGTG<br>GCTACTCTCGAGGATGAGACAGAA<br>GGACCAGGAGTTTGAGGGTAGC<br>TCAGATATGCAATAAGTTCAAGGC<br>CAACCTGTACTATGTTTAAATAGTA<br>AGACAGCATCTCGATAAAATA<br>ATAAAACTAAAGTCTCAACAAAAT<br>AAAAGCTTTCACCTATTAAGGTGCT<br>TGCTTGTCCTTGGAGTCCCCC<br>AAGAGTAACTGCTATGTTAATATCT<br>GTAGAAAGATGTTTATATTTGACT<br>G | https://www.ncbi.nlm.nih.gov/nuccore/NM_028016.3 |
| Sox2 | CAAGCGGCTGCGCGCTCTGCACAT<br>GAAGGAGCACCCGGATTATAAATA<br>CCGGCCGCGGCGGAAAACCAAGA<br>CGCTCATGAAGAAGGATAAGTACA | GCAAGCAACTTTTGTACAGTATTTA<br>TCGAGATAAACATGGCAATCAAAT<br>GTCCATTGTTTATAAGCTGAGAATT<br>TGCCAATATTTTTCGAGGAAAGGG | https://www.ncbi.nlm.nih.gov/nuccore/NM_011443.4 |

| | | | |
|---|---|---|---|
| | CGCTTCCCGGAGGCTTGCTGGCCC<br>CCGGCGGGAACAGCATGGCGAGC<br>GGGGTTGGGGTGGGCGCCGGCCT<br>GGGTGCGGGCGTGAACCAGCGCA<br>TGGACAGCTACGCGCACATGAACG<br>GCTGGAGCAACGGCAGCTACAGC<br>ATGATGCAGGAGCAGCTGGGCTAC<br>CCGCAGCACCCGGGCCTCAACGCT<br>CACGGCGCGGCACAGATGCAACC<br>GATGCACCGCTACGACGTCAGCGC<br>CCTGCAGTACAACTCCATGACCAG<br>CTCGCAGACCTACATGAACGGCTC<br>GCCCACCTACAGCATGTCCTACTCG<br>CAGCAGGGCACCCCCGGTATGGCG<br>C | TTCTTGCTGGGTTTTGATTCTGCAG<br>CTTAAATTTAGGACCGTTACAAACA<br>AGGAAGGAGTTTATTCGGATTTGA<br>ACATTTTAGTTTTAAAATTGTACAA<br>AAGGAAAACATGAGAGCAAGTACT<br>GGCAAGACCGTTTTCGTGGTCTTG<br>TTTAAGGCAAACGTTCTAGATTGTA<br>CTAAATTTTTAACTTACTGTTAAAG<br>GCAAAAAAAAAATGTCCATGCAGG<br>TTGATATCGTTGGTAATTTATAATA<br>GCTTTTGTTCAATCCT | |
| Klf4 | CCTCCTAGCCCGGAGGGAGACCGA<br>GGAGTTCAACGACCTCCTGGACCT<br>AGACTTTATCCTTTCCAACTCGCTA<br>ACCCACCAGGAATCGGTGGCCGCC<br>ACCGTGACCACCTCGGCGTCAGCT<br>TCATCCTCGTCTTCCCCGGCGAGCA<br>GCGGCCCTGCCAGCGCGCCCTCCA<br>CCTGCAGCTTCAGCTATCCGATCCG<br>GGCCGGGGGTGACCCGGGCGTGG<br>CTGCCAGCAACACAGGTGGAGGG<br>CTCCTCTACAGCCGAGAATCTGCG<br>CCACCTCCCACGGCCCCCTTCAACC<br>TGGCGGACATCAATGACGTGAGCC<br>CCTCGGGCGGCTTCGTGGCTGAGC<br>TCCTGCGGCCGGAGTTGGACCCAG<br>TATACATTCCGCCACAGCAGCCTCA<br>GCCGCCAGGTGGCGGGCTGATGG<br>GCAAGTT | CTAACCTTTCACACTGTCTTCCCAC<br>GAGGGGAGGAGCCCAGCTGGCAA<br>GCGCTACAATCATGGTCAAGTTCC<br>CAGCAAGTCAGCTTGTGAATGGAT<br>AATCAGGAGAAAGGAAGAGTTCA<br>AGAGACAAAACAGAAATACTAAAA<br>ACAAACAAACAAAAAAACAAACAA<br>AAAAAACAAGAAAAAAAAATCACA<br>GAACAGATGGGGTCTGATACTGGA<br>TGGATCTTCTATCATTCCAATACCA<br>AATCCAACTTGAACATGCCCGGAC<br>TTACAAAATGCCAAGGGGTGACTG<br>GAAGTTTGTGGATATCAGGGTATA<br>CACTAAATCAGTGAGCTTGGGGGG<br>AGGGAAGACCAGGATTCCCTTGAA<br>TTGTGTTTCGATGATGCAATACACA<br>CGTAAAGATCACCTTGTATGCTC | https://www.ncbi.nlm.nih.gov/nuccore/NM_010637.3 |
| Shh | AAAAGCTGACCCCTTTAGCCTACA<br>AGCAGTTTATTCCCAA<br>CGTAGCCGAGAAGACCCTAGGGG<br>CCAGCGGCAGATATGAAGGGAAG<br>ATCACAAGAAACTCCGAACGATTT<br>AAGGAACTCACCCCCAATTACAAC<br>CCCGACATCATATTTAAGGATGAG<br>GAAAACACGGGAGCAGACCGGC<br>TGATGACTCAGAGGTGCAAAGACA<br>AGTTAAATGCCTTGGCCATCTCTGT<br>GATGAACCAGTGGCCTGGAGT<br>GAAGCTGCGAGTGACCGAGGGCT<br>GGGATGAGGACGGCCATCATTCAG<br>AGGAGTCTCTACACTATGAGGGT<br>CGAGCAGTGGACATCACCACGTCC<br>GACCGGGACCGCAGCAAGTACGG<br>CATGCTGGCTCGCCTGGCTGTGG<br>AAGCAGGTTTCGACTGGGTCTACT<br>ATGAATCCAAAGCTCACATCCACT<br>G | AGCGACTGCGAAATAAGGAACTGA<br>TGGGAAAGCGCACGGAAGGAGAC<br>TTTTAATTATAAG<br>AATAATTCATAATAATAATAATAAT<br>GATAATAATAATAATAATAAGTAG<br>GGCAGTCCAAAGTAGACTATA<br>AGGAAGCAAAAACCCCGGGGAGT<br>TCTGTTGTTATGTTTAGTTTATATAT<br>TTTTTTGAAATTTTTCGTTAT<br>TGTCTTATATGGGTTGTTTTTCTCCT<br>CTCCTGGCTATTTATTTGTTTCGTAT<br>GAATAGATGTTTTAAAAA<br>TATGAACGGACCTTCAAGAGCCTT<br>AACTAGTTTGTGTCTTGGATAATTT<br>ATTATTGTGTGAACTGTACTC<br>ACAGTGAGGGAAAGATTATTTTGT<br>GAGGCCAAGCAACCTGCTGAAAGT<br>CTATTTTTCTACATGTCCCTTG<br>TCCTGCGTTTCAGAAGGCAAACCT<br>C | https://www.ncbi.nlm.nih.gov/nuccore/NM_009170.3 |

| | | | |
|---|---|---|---|
| Netrin1 | GCTG<br>CAGATTCAACATGGAGCTCTATAA<br>GCTATCAGGGCGCAAGAGCGGGG<br>GAGTCTGTCTCAACTGCCGCCAC<br>AACACTGCGGGCCGCCACTGCCAC<br>TACTGCAAGGAGGGCTTCTACCGA<br>GACATGGGCAAGCCTATCACCC<br>ACCGGAAGGCTTGCAAAGCCTGTG<br>ATTGCCACCCAGTGGGTGCTGCTG<br>GCAAGACCTGCAATCAAACCAC<br>TGGCCAATGTCCCTGCAAGGACGG<br>CGTGACGGGCATCACCTGCAACCG<br>ATGTGCCAAAGGCTACCAGCAG<br>AGCCGCTCCCCCATCGCCCCTTGCA<br>TCAAGATTCCTGTGGCGCCACCCA<br>CCACTGCAGCCAGCAGCGTGG<br>AGGAACCGGAAGACTGTGACTCCT<br>ATTGCAAGGCCTCCAAAGGCAAGC<br>TGAAGATGAACATGAAGAAATA<br>CTGCAGGAAGGACTATGCTGTCCA<br>G | GAGAGTGG<br>GCAAGATAGGCTCACTAATGGGCC<br>GTGGTTCACAGACAGATATTCCTG<br>TGGACCAGAGCCATGCCATACC<br>CCAGGGGTATCAAAATTGTCTTTGT<br>GGGGCCTTGCCCCTGCCAAAAACT<br>AAGCCAGCCTCACCTCTTGTG<br>TCAGCGGACTTCCTTCCCCTTTCCC<br>TGATCTGGGTACACCCCCCCGGCC<br>TCCCTCCTGTGTCACCGATTC<br>TGCTCACACAGAATTGTAAATGTTT<br>AGTTGTGACCATGACATATTGTTTG<br>GGCCAGTGTTCCTTTCCAAT<br>GCATACTAATATATTATGGTTATTA<br>TATATGAATATATTTAATGACATGG<br>AGAAAGTTGTGGATTTTCTT<br>TCTTTTTCTTTCTTTTTTTTTTTAAAG<br>TTTTTTTGTTGTTAGAGTTGTAATG<br>GACCCAGACGGAACTTGT<br>AACGTGGGCCCTACAT | https://www.ncbi.nlm.nih.gov/nuccore/NM_008744.2 |
| DCC | TTCTTCAGCGCAGCAGAGGAAGAA<br>ACGGGCCACACACAGTGTGAGCAA<br>GAGGAAGGGCAGTCAGAA<br>GGACCTCCGGCCTCCCGATCTTTG<br>GATACATCATGAAGAAATGGAAAT<br>GAAAAATATCGAGAAGCCTACG<br>GGAACCGACCCTGCAGGAAGAGA<br>CTCCCCCATCCAGAGCTGCCAAGA<br>TCTCACACCAGTCAGCCATAGCC<br>AGTCAGAAACCCAGATGGGAAGC<br>AAAAGTGCCTCTCATTCAGGTCAG<br>GACACTGAGGACGCAGGCAGCTC<br>CATGTCCACTTTGGAACGGTCCCTG<br>GCAGCACGCCGGGCCACCAGGGC<br>CAAGCTCATGATTCCCATGGAG<br>GCCCAGTCCAGTAATCCTGCTGTTG<br>TGAGTGCCATCCCTGTACCAACACT<br>AGAAAGTGCCCAGTATCCAG<br>GAATCCTCCCATCTCCCACATGTGG<br>ATACCCGCATCCACAGTTCACTCTC<br>CGGCCAGTACCATTCCCAAC<br>GCTGTCTGTGGACCGAGGTTTTGG<br>AGCAGGAAGAACTCAGTCTGTGAG<br>TGAAGGACCAACCACGCAACAG<br>CAACCCATGCTGCCCCCAGCTCAG<br>CCCGAACATCCGAGCAGTGAAGAA<br>GCACCCAGTAGAACCATCCCCA<br>CAGCCTGTGTGCGGCCAACTCATC<br>CACTCCGCAGCTTTGCTAACCCATT<br>ACTACCTCCACCCATGAGTGC<br>AATAGAACCA | TCTGGACCATTCGGTTAAGGAACA<br>TCTGAAGCACTT<br>CAGAAGAAAACAGTATCCCATCCT<br>ATATGGACATCAGGAGTATACGAT<br>GATGGTCATGAGGTCCCGAAGA<br>TGCATCTTCACTGCAAAGTCAAGG<br>CTGAAGCTGCTAGCATAGTTCTGG<br>GTCTTTGTCACTGCAGGGACCA<br>TGGTGTCATAACTAATGCCTACATT<br>TTCCTTCAGTGAGACCTTTCATTTTC<br>TGTGTAGGTCTAGTCTCAC<br>AAGTGCATTATTTCAGCCTGTACCA<br>TGCCATAACAAATCAATACAAACTC<br>ACTATTTTTTTCAGCTACAG<br>AGCATCCATGGTAACTCTCTCACTC<br>CATAGCACCAAAGGATTGGATATT<br>TTCCTGACAGCATAAAAGAAA<br>ATAAGTGAATAAGCAAAAGGCAGT<br>GAATGCT | https://www.ncbi.nlm.nih.gov/nuccore/NM_007831.3 |
| Unc5b | GTACTCGGACCTGCACCAATCCAG<br>CCCCACTCAATGGAG<br>GCGCCTTCTGTGAGGGACAGGCCT<br>TCCAGAAGACAGCTTGCACCACCG | ATGTGTGTGTGGGTGTGGTCAATA<br>AGAGACT<br>GCACAGGGAAAGGCTTTGCTATGT<br>CAAGGCCTCAGTTTCCCCAGCTGT | https://www.ncbi.nlm.nih.gov/nuccore/NM_029770.3 |

| | | | |
|---|---|---|---|
| | TGTGCCCAGTGGATGGAGCGTG<br>GACCGAGTGGAGCAAGTGGTCTG<br>CCTGCAGCACAGAGTGTGCGCACT<br>GGCGCAGCCGCGAGTGCATGGCA<br>CCGCCACCCCAGAACGGAGGCCGT<br>GACTGCAGCGGGACGCTACTTGAC<br>TCCAAGAACTGCACTGATGGGC<br>TGTGCGTGCTGAATCAGAGAACTC<br>TAAACGACCCTAAAAGCCACCCCCT<br>GGAGACATCGGGAGATGTGGC<br>ACTGTACGCAGGCCTTGTGGTGGC<br>CGTCTTTGTGGTGGTAGCGGTTCTC<br>AT | GCAGGGAGCTGAGACAGGGTTT<br>TAGCCATGGCCCTGAGTTCTAAAG<br>TCACAAGCAGAAGCTGGGGCCTG<br>GTGGCCTTCGTCCTCTCCGGAGC<br>CCCCTGCAGGTGCCCTTCTGTGGCC<br>TCCTCTGGCTTTGGAGGGTCCAGC<br>TGTGGTTCTGGGAGACTGTTC<br>AGGTCTGTTTGGTGATTCCTGGTCT<br>CGTGATGGCCAAAACTCTTACTGA<br>GAGGGAGGGGGTGTTCCAGGG<br>AGGGCTCTCTCCCTGGGTGAGGCT<br>GGGTGGCTCTGTGTCTAAGCTGGC<br>ACTAGCCGGTCTGCACCCCTGT<br>GCCTCTAGCCATCATTTTACCCGCT<br>GATCTCCCTACTGGGGTCAGGAGA<br>TATGAAGGTAACTAAGTAGAC<br>AGCTGTATACCCACTCTTGACCACA<br>GCGCATTCTGGGATGTTCCATGGG<br>GCCAGTGGAAGGGATGAATTC<br>TTGTCCCTTTCAGCCCACGGGCAGT<br>GACAGGGACTAGGATCACTGGTAT<br>TGATCCTATTGAGATGCTGAC<br>TTCCCTGAACTGCCCTTGACCAGCC<br>TGAAAGCTTTGAAAGTTGGGGCCT<br>CTGTGAGGAGGCAGGTTTCAT<br>GGCCACAGACCCAGCAAGCATCTC<br>TGCCCTGGTTTTCCT | |
| Unc5d | CGGAGTACCATGGCAAGA<br>ATCACTCGGGGACTTTCCCCCATG<br>GAAACAACCGCGGATTCAGTACAA<br>TACATCCCAGAAACAAAACGCC<br>GTACATCCAAAATCTGTCATCACTG<br>CCGACAAGGACAGAGCTGAGGAC<br>AACTGGTGTCTTTGGCCATTTA<br>GGGGGACGCTTAGTAATGCCAAAT<br>ACAGGGGTGAGTCTACTCATACCA<br>CATGGTGCCATCCCAGAGGAGA<br>ATTCTTGGGAGATTTATATGTCAAT<br>CAACCAAGGTGAACCGAGCCTGCA<br>GTCAGATGGATCTGAGGTTCT<br>CCTGAGTCCTGAAGTCACCTGTGG<br>GCCTCCAGATATGCTTGTCACAACT<br>CCCTTTGCGCTGACCATCCCT<br>CACTGTGCAGACGTCAGTTCAGAG<br>CACTGGA | CACTGTGACCCCAAGCTTTTGCATG<br>ACTTTCACCTGTCACAGCTCAAATG<br>CCCATCCTGGAAGAGGTTT<br>CCACATTACTCTGGGCGCTAAATTC<br>GGAAGTGTCACTGTCCTAGTGCAA<br>GATAAAACAAAAATTAAAAGT<br>GTCCTTAAATACTTAAAAACATAAA<br>TTTTTTACAAAAACAATCCTACTCA<br>ATGCCAAATTTCTCAGTGCT<br>TGGAGACATTAAATCTATACTTTAA<br>ATATGAGTTTAAATTTTGCCACAAA<br>AATGGTGAGATCCACAGGTT<br>TGCTAGACTACAGCAGCATTACGT<br>TAATTTACAGTCTCTACCACTCCAC<br>ATACACAGTCTGAGTAGTTTA<br>CATTTGCATTCTTTAATCCAAAAGG<br>AAACTTGTTCTGTGGTTTTAAATGC<br>TACCTTCATTCACCAGTGCG<br>GGTAGAC | https://www.ncbi.nlm.nih.gov/nuccore/NM_153135.3 |
| Hnrnpa0 | CCTCAATGTGCAGACGAGTGAGTC<br>GGGGCTGCGCGGCCACTTCGAGG<br>CCTTCGGGACGCTGACGGACTG<br>CGTGGTGGTGGTGAACCCCCAGAC<br>GAAGCGCTCCCGCTGCTTCGGCTT<br>CGTGACCTACTCGAACGTGGAG<br>GAGGCGGATGCCGCCATGGCCGC<br>GTCGCCGCACGCGGTGGACGGCA<br>ACACGGTGGAGCTGAAGCGCGCC<br>G | ATAGCTGCAAGACGGTTGGTTAA<br>GAAAAACCCAACTTACTTGTTCAG<br>GACAATTTCATCAGATTTAGCAGTT<br>TGCCATCAGAATTCTGCAGAC<br>TTGAAAGGTCCACATTTACTTGTAG<br>CTGAGACGGGCTTCCCGGGAGTGT<br>AGTTAACAGCCCGCATCAGTC<br>ACTTTTTCTACGTGAGATTTGAACA<br>TGTAAACTGCTATGCATAGCTTGG<br>GCAGTAGCCCCAAACTGCTCC | https://www.ncbi.nlm.nih.gov/nuccore/NM_029872.1 |

| | | | |
|---|---|---|---|
| | TGTCGCGGGAGGATTCGGCGCGG CCCGGGGCGCACGCCAAGGTGAA GAAGCTGTTCGTGGGCGGCCTCAA GGGCGACGTGGCGGAGGGCGACC TGATCGAGCACTTCTCGCAGTTCG GCGCGGTGGAGAAGGCGGAGATC ATTGCCGACAAGCAGTCGGGCAAG AAGCGCGGCTTCGGCTTCGTCTAC TTC | GGTAGCTACTGAAGCAGCTGTGTG TACTGGGGTTTTAGGTGCAAGTTG TAAAACAAAATATCTGTATTCT GCTTGGTTAACGTGTATTTGTAGCC CTTCATGCAATAGAATTCAAGTTGT TGTTTATAAAAAAAAAACAA AAACTGATAATAGAAGAAAATGCC TTCCTGGA | |
| Oct4 | GTGATGGGTC AGCAGGGCTG GAGCCGGGCT GGGTGGATCC TCGAACCTGGCTAAGCTTCCAAGG GCCTCC AGGTGGGCCT GGAATCGGAC CAGGCTCAGA GGTATTGGGGATCTCCCCATGTCC GCCCGC ATACGAGTTC TGCGGAGGGA TGGCATACTG TGGACCTCAGGTTGGACTGGGCCT AGTCCC CCAAGTTGGC GTGGAGACTT TGCAGCCTGA GGGCCAGGCAGGAGCACGAGTGG AAAGCAA CTCAGAGGGA ACCTCCTCTG AGCCCTGTGC CGACCGCCCCAATGCCGTGAAGTT GGAGAA GGTGGAACCA ACTCCCGAGG AGTCCCAGGA CATGAAAGCCCTGCAGAAGGAGCT AGAACAGTTTGCCAAGCTGCTGAA GC AGAAGAGGAT CACCTTGGGG TACACCCAGGCCGACGTGGG | CCTGGGGATGCTGTGAGCCA AGGCAAGGGA GGTAGACAAG AGAACCTGGAGCTTTGGGGTTAAA TTCTTT TACTGAGGAG GGATTAAAAG CACAACAGGG GTGGGGGGTGGGATGGGGAA AGAAGCTCAG TGATGCTGTT GATCAGGAGC CTGGCCTGTC TGTCACTCATCATTTTGTTCTTAAAT AAAGACTGGGACACACAGTAGATA | https://www.ncbi.nlm.nih.gov/nuccore/NM_013633.3 |
